# CLIP-Seq Analysis Enables the Design of Ribosomal RNA Bait Oligonucleotides That Protect Against *C9ORF72* ALS/FTD-Associated Poly-GR Pathophysiology

**DOI:** 10.1101/2022.12.30.522259

**Authors:** Juan A. Ortega, Ivan R. Sasselli, Marco Boccitto, Andrew C. Fleming, Tyler R. Fortuna, Yichen Li, Kohei Sato, Tristan D. Clemons, Elizabeth L. Daley, Thao P. Nguyen, Eric N. Anderson, Justin K. Ichida, Udai B. Pandey, Sandra Wolin, Samuel I. Stupp, Evangelos Kiskinis

## Abstract

Amyotrophic lateral sclerosis and frontotemporal dementia patients with a hexanucleotide repeat expansion in *C9ORF72* (C9-HRE) accumulate poly-GR and poly-PR aggregates. The pathogenicity of these arginine-rich dipeptide repeats (R-DPRs) is thought to be driven by their propensity to bind to low complexity domains of multivalent proteins. However, the ability of R-DPRs to bind native RNA and the significance of this interaction remains unclear. We used computational and experimental approaches to characterize the physicochemical properties of R-DPRs and their interaction with RNA. We find that poly-GR predominantly binds ribosomal RNA (rRNA) in cells and exhibits an interaction that is predicted to be energetically stronger than that for associated ribosomal proteins. Critically, modified rRNA “bait” oligonucleotides restore poly-GR-associated ribosomal deficits in cells and ameliorate poly-GR toxicity in patient neurons and *Drosophila* models. Our work strengthens the hypothesis that ribosomal function is impaired by R-DPRs, highlights a role for direct rRNA binding in mediating ribosomal disfunction, and presents a strategy for protecting against C9-HRE pathophysiological mechanisms.

## INTRODUCTION

Amyotrophic lateral sclerosis (ALS) and frontotemporal dementia (FTD) are two progressive and untreatable neurodegenerative diseases with overlapping genetic abnormalities but divergent clinical presentations. The major genetic cause of both diseases is a heterozygous intronic hexanucleotide (GGGGCC)_n_ repeat expansion in the *C9ORF72* gene (C9-HRE) (DeJesus-Hernandez et al., 2011; Renton et al., 2011). The C9-HRE causes a reduction in C9ORF72 protein and produces neurotoxic RNA (Donnelly et al., 2013; McEachin et al., 2020). Both the sense and antisense GGGGCC expansion can be transcribed and translated into 5 distinct dipeptide repeat (DPR) proteins (glycine-proline (GP), glycine-alanine (GA), glycine-arginine (GR), proline-arginine (PR), and proline-alanine (PA) (Ash et al., 2013; Gendron et al., 2013; Mori et al., 2013; Zu et al., 2013). Although the relative pathogenic contribution of these mechanisms to the disease remains unclear, multiple studies have demonstrated that C9-DPRs, and especially the arginine-rich GR and PR, are particularly toxic (reviewed by (Balendra and Isaacs, 2018; Freibaum and Taylor, 2017; Haeusler et al., 2016). Expression of GR and PR results in cellular toxicity *in vitro*, as well as stark neurodegeneration in *Drosophila* models, and neurodegeneration and behavioral phenotypes in mouse models *in vivo* (Choi et al., 2019; Freibaum et al., 2015; Hao et al., 2019; Lee et al., 2016; Mizielinska et al., 2014; Tao et al., 2015; Wen et al., 2014; Zhang et al., 2018b; Zhang et al., 2019).

The pathogenicity of poly-GR and poly-PR is thought to be primarily driven by the propensity of their arginine-rich core sequence to bind to low complexity domains (LCD) of multivalent proteins by ionic and cation-pi contacts. Indeed, a number of mass-spectrometry (MS)-based interaction assays have demonstrated that R-DPRs heterogeneously bind to LCD-containing proteins that are associated with multiple cellular functions including RNA metabolism and nucleocytoplasmic (N/C) trafficking (Hartmann et al., 2018; Jovicic et al., 2015; Kanekura et al., 2016; Kwon et al., 2014; Lee et al., 2016; Lin et al., 2016; Radwan et al., 2020; Tao et al., 2015; White et al., 2019; Yin et al., 2017; Zhang et al., 2018a). However, while R-DPRs have been shown to bind RNA *in vitro* (Boeynaems et al., 2017; Kanekura et al., 2016; Lee et al., 2016; Lin et al., 2016; White et al., 2019), nothing is known about their native interactions with RNA in living cells, and the functional consequences of these interactions. To address this critical gap in knowledge, we combined empirical and bioinformatic tools to decipher how the polar nature and the differential secondary structure of R-DPRs determine their interaction with RNA. Cross-linking immunoprecipitation followed by high throughput sequencing, supported by targeted Northern blot and quantitative PCR experiments, identified various ribosomal RNA (rRNA) species as direct targets of poly-GR in living cells. This interaction, which occurs in both the nucleus and cytoplasm, disrupts ribosomal homeostasis. Based on these findings, we designed a modified rRNA oligonucleotide that acted as a bait and inhibited poly-GR associated ribosomal defects. The rRNA bait ameliorated the toxicity of poly-GR in iPSC-derived motor neurons and cellular models *in vitro* and *Drosophila* models *in vivo*. Our work strengthens the hypothesis that ribosomal function is severely impaired by R-DPRs and suggests that a sequence specific rRNA molecule can abrogate the toxic effects of poly-GR.

## RESULTS

### Characterization of the physicochemical features of C9 arginine (R)-DPRs

To interrogate the role of RNA in DPR pathophysiology we first characterized the structural and chemical features of (R)-DPRs. We specifically examined the secondary structure of the R-rich, highly toxic poly-GR and poly-PR by all atoms molecular dynamics (MD) simulations using the CHARMM force field (Figure 1A-B). As a control, we looked at the non-toxic poly-GP DPR. We modeled DPRs with 15 repeats, which represents the maximum length of peptides we could synthesize with high purity and corresponds to half of the minimum number of hexanucleotide (GGGGCC)n repeats that is considered to be neurotoxic (n=30). While both (GP)_15_ and (GR)_15_ showed a highly coiled conformation, (PR)_15_ displayed a more stretched structure (Figures 1A-B and S1A). The differential physicochemical properties of proline and glycine likely impact the differential conformation of DPRs with proline favoring a rigid backbone, and glycine conferring higher flexibility (Jafarinia et al., 2020). Along these lines, the calculation of torsional angles in the different DPR residues by Ramachandran plots showed that (PR)_15_ has a high enrichment in β-sheet conformations, while (GR)_15_, and (GP)_15_ displayed a more diverse profile of different secondary conformations in their structures (Figure 1C). This correlates with a higher extended conformation observed in (PR)_15_, while (GP)_15_ and (GR)_15_ show more convoluted structures (Figure 1B). To validate the results obtained by MD simulations we synthesized highly pure DPRs with 15 repeats (Figure S1B) and characterized them by circular dichroism (CD) (Micsonai et al., 2015). Using spectra of proteins and peptides with highly pure secondary structure contributions as a reference (Figure 1D, right), we obtained qualitative information of the major secondary structures in C9-DPRs. We found that while (PR)_15_ showed a β-sheet spectrum with the characteristic negative peak over 210nm, (GP)_15_ showed two negative peaks, typical of an α-helix, and (GR)_15_ displayed a random coiled spectrum (Figures 1D). Additionally, we used the BeStSel tool (Micsonai et al., 2018) to analyze the secondary structure ratios in the three distinct DPR proteins, and found major contributions of helix, sheet and coil conformations in (GP)_15_, (PR)_15_, and (GR)_15_ respectively (Figure 1E).

**Figure 1.**
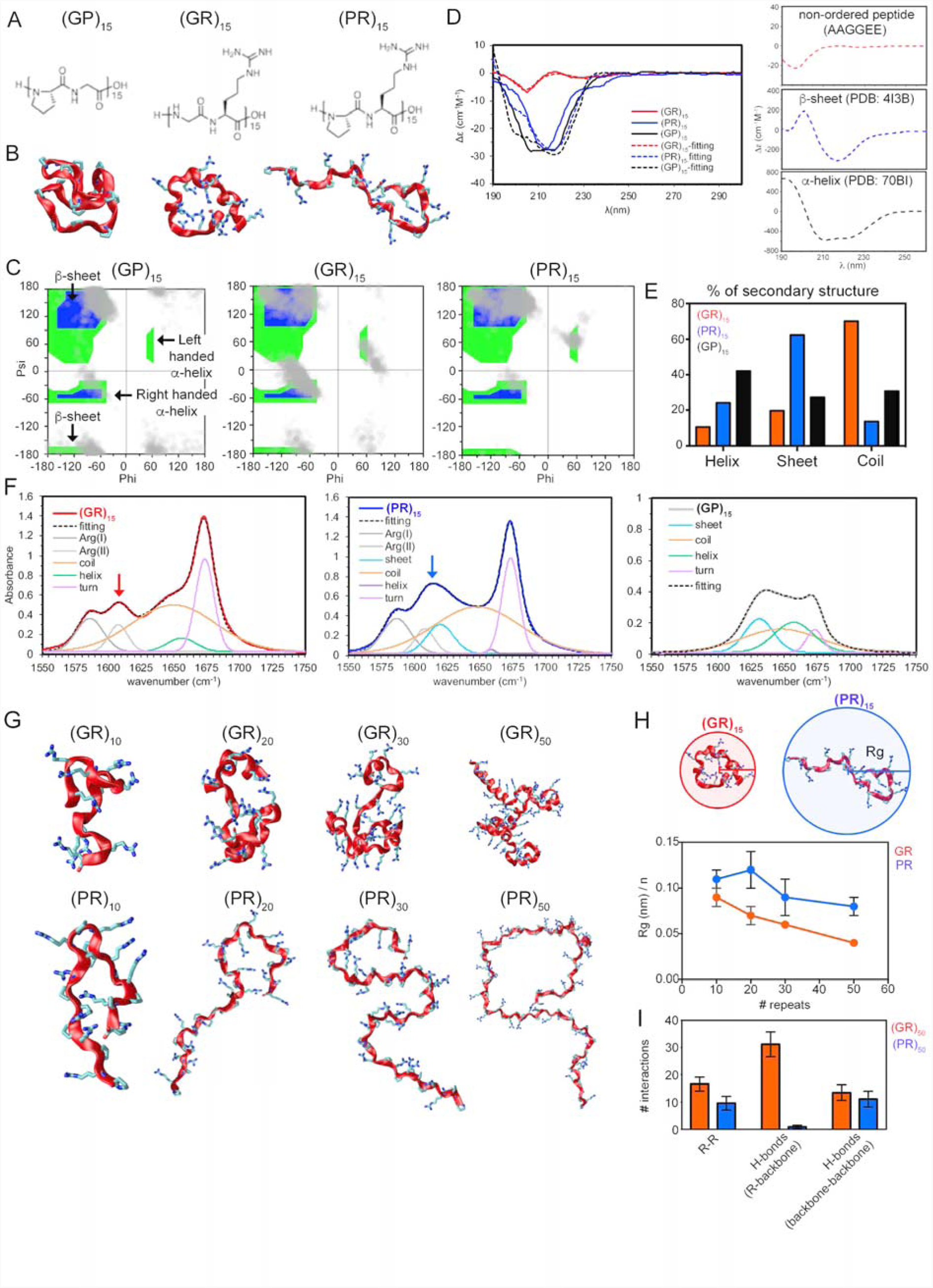
Computational and empirical characterization of the physicochemical features of C9 R-DPRs. **A**. Scheme showing the of (GP)_15_, (GR)_15_ and (PR)_15_ chemical structures. **B**. (GP)_15_, (GR)_15_, and (PR)_15_ structures after all-atoms molecular dynamic simulations. Backbone shows the secondary structure and side chain hydrogens are omitted for clarity. **C**. Ramachandran plots mapping the frequencies of secondary structures (gray dots) present within (GP)_15_, (GR)_15_ and (PR)_15_. Green and blue areas identify different types of dihedral angles associated to β-sheets, left- and right-handed α-helices as shown in left panel. **D**. Left: Line graph showing the secondary structure traces of 1mM (GP)_15_, (GR)_15_ and (PR)_15_ in 10mM HEPES, 10mM NaCl pH7.2, analyzed by circular dichroism (CD). Dashed lines showed the fitting convolution of (GP)_15_, (GR)_15_ and (PR)_15_-CD spectra. Right: Line graphs showing the CD traces of non-ordered, β-sheet rich (PDB: 4I3B) and α-helix rich (PDB: 7OBI) peptides used as a reference. **E**. Bar plot depicting the percentage of secondary structures in (GP)_15_, (GR)_15_ and (PR)_15_ based on our CD analysis. **F**. Line graph showing FTIR spectra of (GP)_15_, (GR)_15_, and (PR)_15_ in the amide I region. Spectra were deconvoluted to show the secondary structure traces (sheet, coil, helix, and turn) and arginine side chain contributions (Arg I and Arg II). HEPES solution was used as a control. **G**. All-atoms molecular dynamics simulation results of (GR)_n_ (top) and (PR)_n_ (bottom) with different number of repeats (n=10, 20, 30, 50) representing their secondary structures. **H**. Top: schematic representation of the peptide length calculation for (GR)_15_ and (PR)_15_, measured by the radius of the circle that circumscribes each R-DPR. Bottom: bar graph depicting the radius of gyration (Rg) measure in the simulations of (GR)_n_ and (PR)_n_ with different number of repeats (n=10, 20, 30, 50). Values are presented as the mean ± standard deviation (SD). **I**. Bar plot showing the number of R-R and H-bond (within the backbone or between backbone and R residues) interactions in (GR)_50_ and (PR)_50_. Values are presented as the mean ± SD.

**Supplementary Figure 1.**
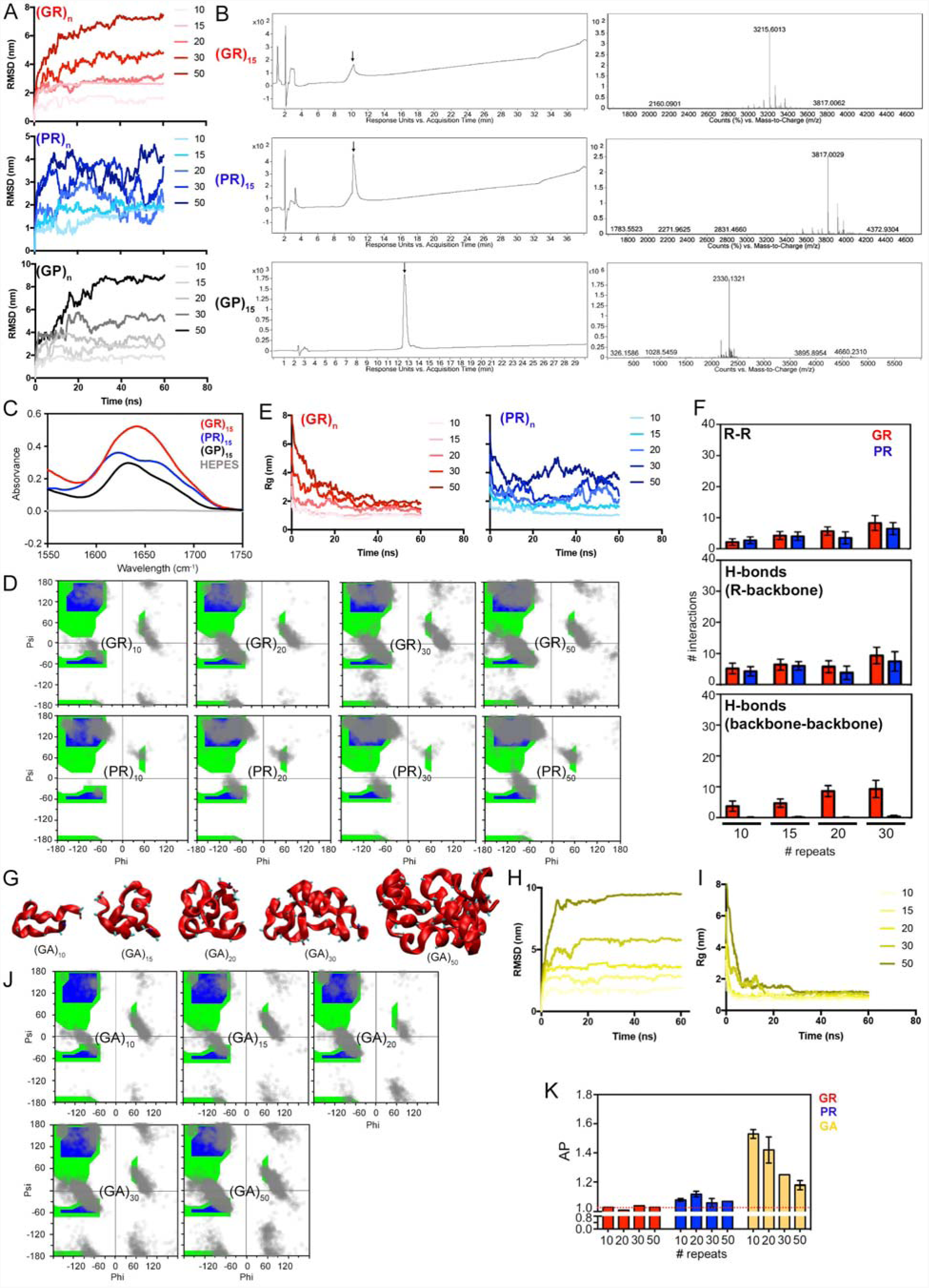
Computational and empirical characterization of the physicochemical features of C9 R-DPRs. **A**. Line graphs showing changes from the initial structure calculated by root-mean-square deviation (RMSD) over time in (GR)_n_ (top), (PR)_n_ (middle), and (GP)_n_ (bottom) with different repeat number (n=10, 15, 20, 30, 50) in molecular dynamics simulations. **B**. LC-MS chromatograms using UV-vis absorbance (left) and ESI-MS spectra (right) obtained from pure fractions of synthesized (GR)_15_, (PR)_15_ and (GP)_15_. **C**. Line graph showing the secondary structure traces of (GP)_15_, (GR)_15_ and (PR)_15_ analyzed by solid FTIR. **D**. Ramachandran plots mapping the secondary structures population present in (GR)_n_ (top) and (PR)_n_ (bottom) with different repeat number (n=10, 15, 20, 30, 50). **E**. Line graphs representing the radius gyration (Rg) changes over time in the molecular dynamics simulations of (GR)_n_ and (PR)_n_ with different repeat number (n=10, 15, 20, 30, 50). **F**. Bar plots showing the number of R-R and H-bond (within the backbone or between backbone and R residues) interactions in (GR)_n_ and (PR)_n_ with different number of repeats (10, 20, 30 and 50). The values are presented as the mean ± SD. **G**. All-atoms molecular dynamics simulation of the secondary structure of (GA)_n_ with different repeat number (n=10, 15, 20, 30, 50). **H**. Line graph showing the changes from the initial structure calculated by RMSD over time in (GA)_n_ with different repeat number (n=10, 15, 20, 30, 50). **I**. Line graphs representing the radius gyration (Rg) changes over time in the molecular dynamics simulations of (GA)_n_ with different repeat number (n=10, 15, 20, 30, 50). **J.** Ramachandran plots mapping the secondary structures population present in (GA)_n_ with different repeat number (n=10, 15, 20, 30, 50). **K**. Bar graphs showing coarse-grained molecular dynamics simulation-based aggregation propensity calculated for (GR)_n_, (PR)_n_ and (GA)_n_ (n=10, 20, 30, 50). Red dashed line indicates the average AP values calculated in poly-GR. Values are presented as the mean ± SD. Raw data of all the simulations presented in this figure can be found in Table S1.

Given the unique nature of R-DPRs and the lack of any similar datasets for comparison, we employed additional techniques to confirm our results. We specifically performed liquid and solid sampling of the C9 DPRs and used Fourier-transform infrared spectroscopy (FTIR), which allows for collecting high-resolution spectral data without a potential impact of solvent composition. We focused on the amide I region, as this vibrational mode is known to be highly sensitive to peptide secondary structure and the shift of the signals are benchmarked for the different types of secondary structures (Barth, 2000). FTIR is not affected by the loss of chirality in glycine-rich sequences, but it can present overlapping signals and interference from other chemical groups, and thus signal deconvolution is essential. We observed contributions from the arginine’s side chain (Arg I and Arg II) at 1587 and 1608 cm^−1^ for (GR)_15_ (Figure 1F) (Barth, 2000), while for (PR)_15_ the latter peak (red arrow in (GR)_15_ signal) is merged with the β-sheet contribution of the amide I, giving a peak at around 1615 cm^−1^ (blue arrow). In contrast, the main amide I contribution of (GR)_15_ is displayed at 1648 cm^−1^, typical of random coil (Figure 1F) (Barth, 2000; Barth and Zscherp, 2002). (GP)_15_ displayed a sum of amide I vibrations with similar intensity suggesting that it exhibits a combination of β-sheet, random coil, α-helix, and turn conformations. The deconvolution shows the strong relevance on the random coil mode to the final spectra and highlights the α-helix contribution in the three peptides, that is typically masked by other peaks. This contribution is especially important in (GP)_15_. While all three DPRs exhibited a signal at 1675 cm^−1^, typical of rigid turns, this was strongly enhanced in R-DPRs (Figure 1F), likely reflecting the additional intramolecular R-R interactions that R-DPRs form as shown in the MD simulations. In solid FTIR analysis, the arginine side chain vibrations disappear, validating the β-sheet nature of the (PR)_15_ amide I vibration around 1620 cm^−1^ (Figure S1C). In contrast, (GP)_15_ and (GR)_15_ exhibited higher amide I frequencies, indicative of a random coiled structure that is more predominant for (GR)_15_ (Figure S1C).

We next asked how the number of repeats would impact the structure of the DPRs using MD simulations. Ramachandran plots of both GR and PR with 10, 20, 30 and 50 repeats showed the same secondary structure patterns, irrespective of length (Figures 1G and S1D). Importantly, the higher flexibility of the poly-GR backbone favors a more folded conformation compared to the poly-PR backbone, which exhibited an extended conformation (Figures 1G and S1E). Accordingly, the higher level of folding observed in poly-GR peptides led to a size-dependent increase in the number of R-R and H-bond interactions within the peptide backbone relative to poly-PR (Figures 1H-I and S1F). We also performed MD simulations to interrogate the intrinsic aggregation propensity (AP) of different size R-DPRs. As a positive control we used poly-GA, because of its well-described tendency to self-aggregate (Chang et al., 2016; Freibaum and Taylor, 2017; Jafarinia et al., 2020) (Figures S1G-J). Although R-DPRs displayed reduced AP profiles compared to poly-GA, the AP values in poly-PR were slightly higher than poly-GR (Figure S1K). Collectively, our computational and empirical analyses indicate that GR and PR exhibit differential secondary structures and aggregation propensities that appear to be only mildly affected by repeat length and could lead to differential pathophysiological effects.

### Characterization of the interaction between C9 R-DPRs and RNA *in vitro*

Previous studies have suggested that R-DPRs can bind to RNA molecules through ionic and cation-pi interactions and undergo liquid-liquid phase separation (LLPS) (Boeynaems et al., 2017; Kanekura et al., 2016; White et al., 2019). Thus, we sought to assess whether the differential physicochemical properties of poly-GR and poly-PR DPRs would impact their binding to RNA molecules. We first incubated total human RNA with synthetic DPRs *in vitro* and measured optical density as an indicator of RNA-DPR interaction (Figures 2A-B and S2A). While (GP)_15_ had no effect, both R-DPRs form complexes with RNA in a concentration-dependent manner (Figure 2A-B). A dose response experiment with increasing concentrations of DPRs showed that the interaction of (GR)_15_ with RNA peaked at 20 μM, while (PR)_15_ peaked at 15 μM, suggesting a stronger potential for PR to bind to RNA (Figure 2B). This is in accordance with its expanded structural conformation (Figures 1 and S1) and by extension, the higher number of cations available for multiple electrostatic or cation-pi interactions with RNAs (Figure 2C). Indeed, calculation of the solvent-accessible surface area (SASA) of both R-DPRs and poly-GP indicated higher values in poly-(PR) than the other DPRs, particularly for the 15 and 30 repeat dipeptides (Figures 2D and S2B). We next interrogated how the concentration of RNA would impact the propensity for R-DPRs to phase separate. We observed that while the gradual increase in RNA concentration of up to 10 μg/μl increased the optical density values, higher concentrations reduced phase separation (Figure 2E), suggesting that the interaction is both RNA and R-DPR concentration-dependent. Finally, we asked whether the R-DPRs differentially interact with distinct types of RNA, including ribosomal (rRNA), transfer (tRNA) and messenger RNA (mRNA) *in vitro*. We found that both R-DPRs interacted more strongly with rRNA than mRNA (Figure 2F), correlating with the fact that in cells they accumulate in nucleoli (Figures S2C-D). Moreover, (GR)_15_ exhibited a stronger interaction with tRNA than mRNA (Figure 2F). Collectively, our findings are in line with previous studies in demonstrating that poly R-DPRs bind RNA molecules *in vitro* (Boeynaems et al., 2017; Kanekura et al., 2016), and suggest that R-DPRs may preferentially recognize structured RNAs, such as tRNA and rRNA.

**Figure 2.**
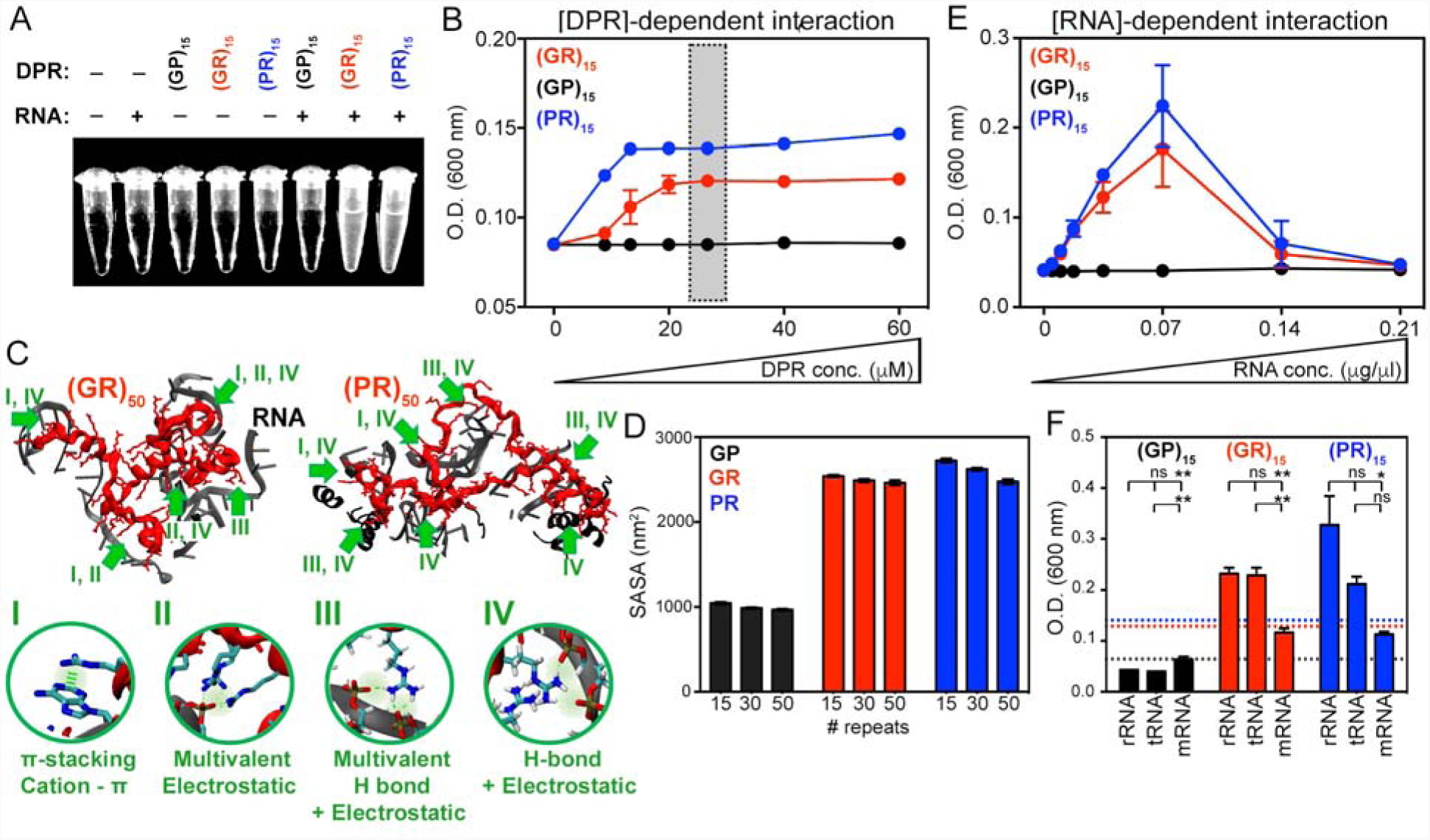
Characterization of the interaction between R-DPRs and RNA *in vitro*. **A**. Photograph of turbidity assays after mixing human total RNA and synthetic (GP)_15_, (GR)_15_ and (PR)_15_. **B**. Line graphs depicting DPR concentration-dependent precipitation of 0.07 μg/μl RNA calculated by optical density. Gray shaded area indicates the DPR concentration utilized in **E**. **C**. MD simulation snapshots showing how differential secondary structures in R-DPRs (red) can influence the number and type of interactions with human ribosomal RNA (rRNA; gray/black). Bottom: Circles showing the different types of interactions that could mediate the binding of (GR)_50_ and (PR)_50_ with rRNA. **D**. Graph bars indicating the Solvent Accessible Solvent Area (SASA) measured from molecular dynamics simulations of (GP)_n_, (GR)_n_, and (PR)_n_ with different number of repeats (n=15, 30, 50). **E**. Line graphs depicting RNA concentration-dependent precipitation of (GR)_15_, (GP)_15_ and (PR)_15_ calculated by optical density measurements. **F**. Bar plots showing turbidity assay measurements utilized to assess the level of interaction of 26.7 μM of (GP)_15_, (GR)_15_ and (PR)_15_ with 5 μg/μl ribosomal (rRNA), transfer (tRNA) and messenger (mRNA) RNA. Dot lines indicate the level of interaction of the distinct DPRs with total human RNA. All values are presented as the mean ± SD; ANOVA *P<0.05; **P<0.001; ns: not significant.

**Supplementary Figure 2.**
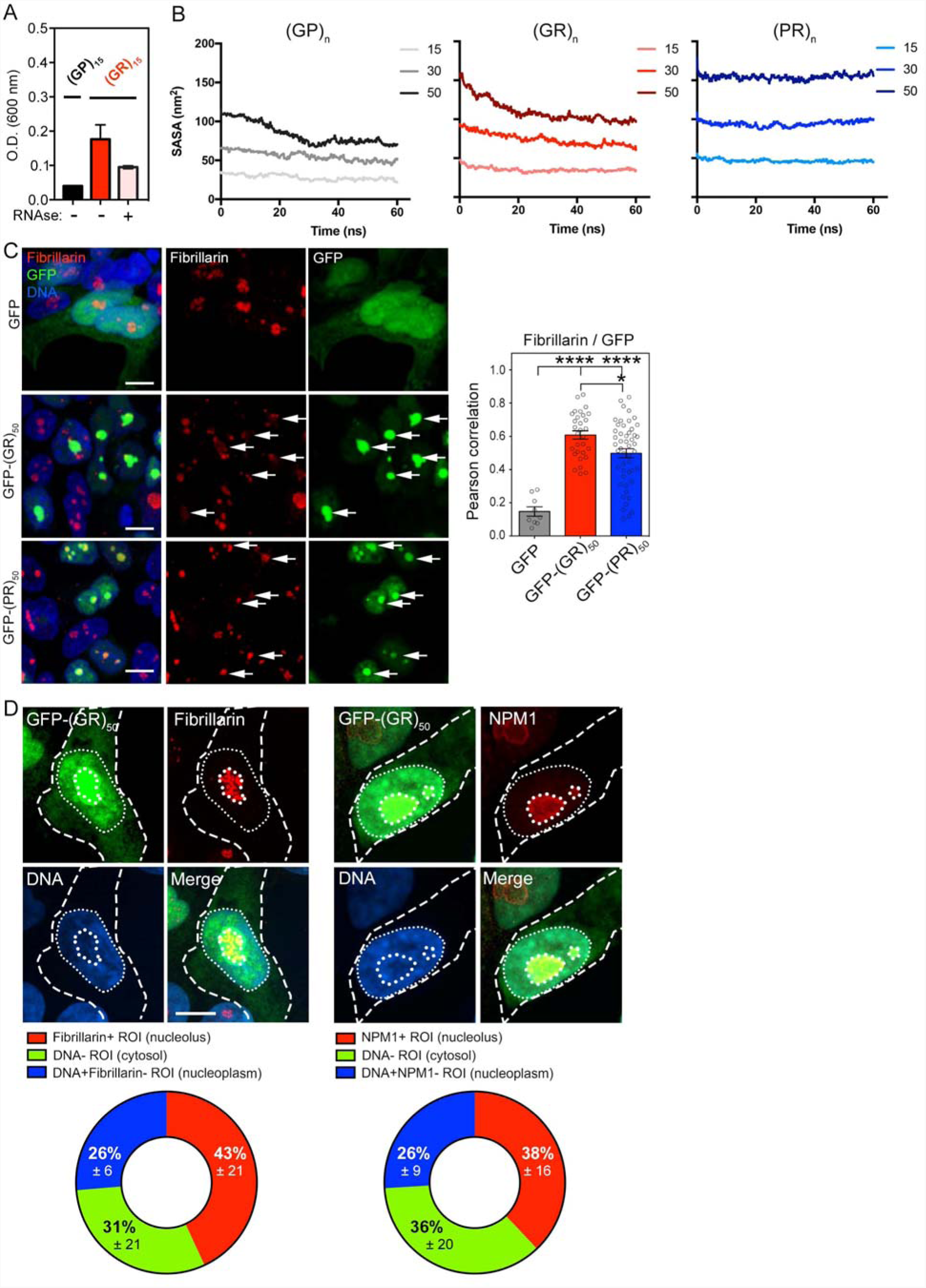
Characterization of the interaction between R-DPRs and RNA *in vitro* and localization of R-DPRs in cells. **A**. Bar plots showing turbidity assay measurements utilized to assess the level of interaction of (GR)_15_ with total human RNA in the absence/presence of RNase. (GP)_15_ was used as a reference as it does not bind to RNA. **B**. Line graphs representing the SASA changes over time in molecular dynamics simulations of (GP)_n_, (GR)_n_, and (PR)_n_ with different number of repeats (n=15, 30, 50). **C**. Left: Confocal images of HEK-293 cells transfected with GFP, GFP-(GR)_50_, or GFP-(PR)_50_ and labeled for the nucleolar marker fibrillarin using immunocytochemistry. Scale bar = 15 μm. Right: Dot plot showing the level of colocalization of fibrillarin with GFP and GFP-fused R-DPRs. Each dot in graphs represents a single cell. **D**. Top: Confocal images of HEK-293 cells transfected with GFP-(GR)_50_ and immunostained for Fibrillarin or NPM1, which label the fibrillar and granular components of the nucleolus respectively. DNA was visualized by staining with Hoechst 33342. Dashed lines indicate cytosolic borders, while thin and thick pointed lines depict nuclear and nucleolar borders, respectively. Bottom: Pie charts displaying the distribution of the GFP-(GR)_50_ signal within the different subcellular compartments (cytosol, nucleoplasm and nucleolus) defined by the selected markers. ROI: region of interest. All values are presented as the mean ± standard error mean (SEM); ANOVA p*<0.05; ***<0.0001.

### Poly-GR binds to ribosomal RNA in cells

We next sought to identify the RNAs bound by poly-GR in live cells. We focused our analysis on poly-GR since GR positive aggregates are more frequently observed in C9-ALS/FTD patient tissue relative to PR aggregates (Gomez-Deza et al., 2015; Mackenzie et al., 2015; Mori et al., 2013), and its abundance has been associated with affected brain areas in patients (Saberi et al., 2018). We used cross-linking immunoprecipitation followed by high-throughput sequencing (CLIP-Seq) (Darnell et al., 2018) to identify targets of poly-GR on a transcriptome-wide scale. In CLIP-Seq, UV light is used to cross-link proteins to RNAs that are in direct contact in live cells. After immunoprecipitating the protein of interest and harsh washing to remove noncovalently associated RNA, cDNA is prepared from the cross-linked RNA and sequenced (Figure 3A). In preliminary experiments, we confirmed that, following transfection into HEK-293 cells, GFP-(GR)_50_ could be crosslinked to RNA (Figures S3A-C). To identify specific targets, we transfected HEK-293 cells with GFP-(GR)_50_, using GFP (Figure 3B) or GFP-(GP)_10_ and untransfected cells (Figure S3D) as controls. Following UV-crosslinking, immunoprecipitation, and cDNA sequencing, we found that GFP-(GR)_50_ interacted predominantly with rRNA. In two biological replicates, 96% and 92% of the GFP-(GR)_50_ CLIP peaks, defined as five or more overlapping sequencing reads at a specific genomic locus, mapped to ribosomal DNA (rDNA) (Figures 3C and S3E; Table S2). Additionally, while 63% and 68% of the CLIP peaks from the control samples also mapped to rDNA, the rRNA-derived peaks were strongly enriched in GFP-(GR)_50_ immunoprecipitates, while tRNAs and Y RNAs that are also highly abundant in cells were de-enriched in GFP-(GR)_50_ samples (Figures 3D and S3F). Close inspection of the RNA45SN1 locus revealed that the GFP-(GR)_50_-derived peaks were not distributed evenly across the rDNA, but rather occurred in specific regions (Figures 3E and S3G).

**Figure 3.**
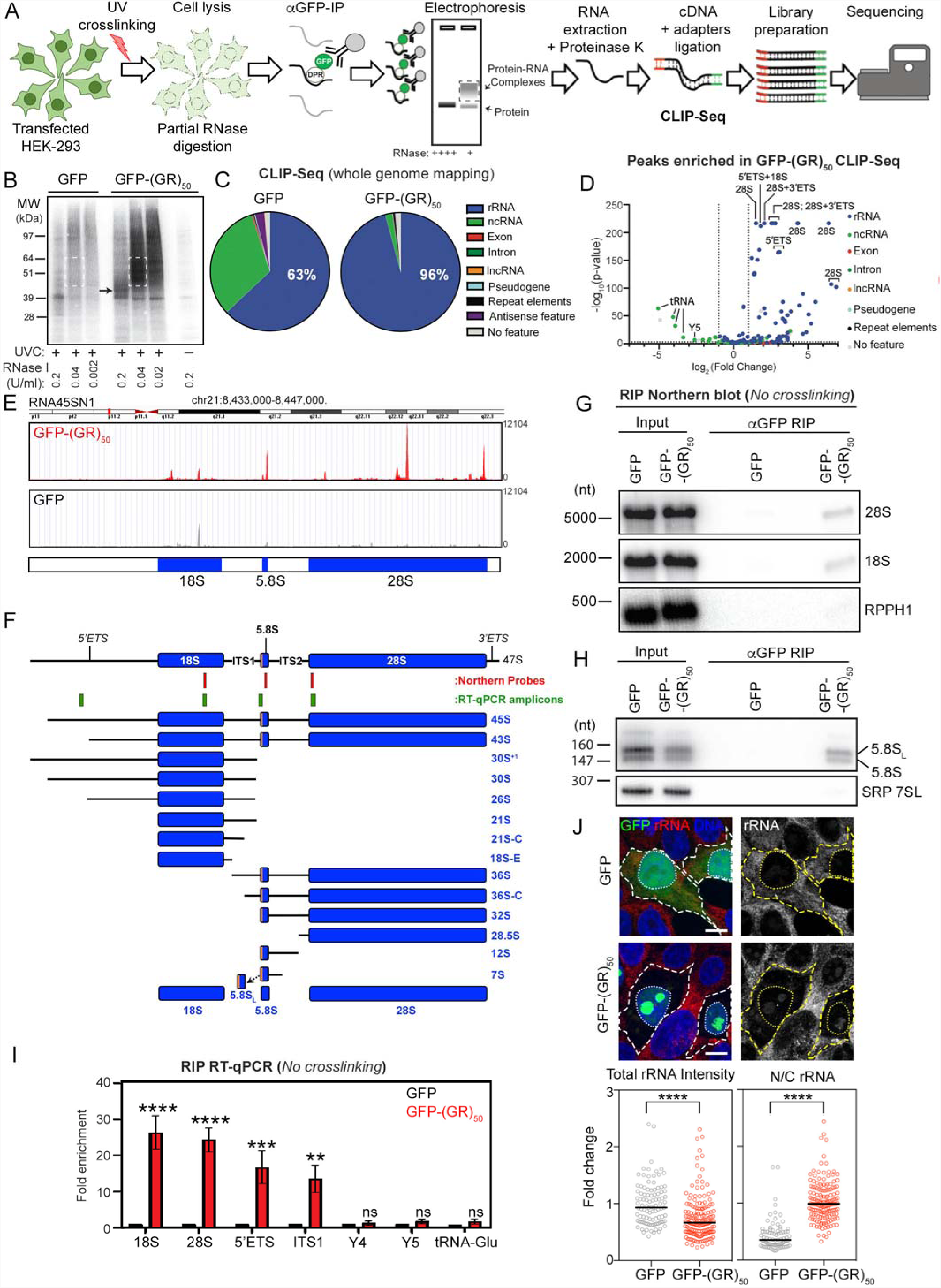
Characterization of the interactions between poly-GR and RNA in live cells. **A**. Schematic of the CLIP-Seq procedure utilized in this study. **B.** Following UV crosslinking, cell lysis, partial ribonuclease (RNase) digestion to generate cloneable fragments and immunoprecipitation with anti-GFP antibodies (as in **A.**). Ribonucleoproteins (RNPs) in precipitates were labeled at the 5’ end with [γ-^32^P]-ATP and fractionated using SDS-PAGE. After transfer to a membrane, RNPs were detected with autoradiography. White dashed lines: RNPs excised for library preparation. Arrow: GFP-(GR)_50_. **C**. Pie charts showing the percentage of CLIP-Seq peaks (replicate #1) found in various RNA classes in GFP *vs.* GFP-(GR)_50_-transfected cells. Peaks were defined as having 5 or more overlapping reads. **D**. Volcano plot showing CLIP-Seq peaks enriched in GFP-(GR)_50_ compared to GFP in CLIP-Seq (replicate #1). Negative log_2_ (fold change) indicates peaks that were more abundant in the GFP control. Some of the most significantly enriched peaks GFP-(GR)_50_-transfected cells are labeled. **E.** Distribution of reads mapping to the RNA45SN1 locus from the GFP-(GR)_50_ (top) and GFP (bottom) samples in CLIP-Seq (replicate #1). **F.** Schematic depicting ribosomal RNA processing and the rRNA precursors that have been detected in human cells (Henras et al., 2015). The positions of probes used for NB as well as the positions of the RT-qPCR amplicons are shown. **G** and **H**. Lysates of HEK-293 cells expressing GFP-(GR)_50_ or GFP were subjected to immunoprecipitation with anti-GFP antibodies. RNAs within immunoprecipitants were fractionated in denaturing agarose (**G**) or polyacrylamide (**H**) gels and the indicated RNAs were detected by NB. The input represents 2% of the lysate used in immunoprecipitation. **I**. After immunoprecipitation, the levels of the indicated RNAs in anti-GFP and anti-GFP-(GR)_50_ immunoprecipitants were measured by RT-qPCR. The fold-enrichment of each RNA relative to the anti-GFP control is shown. (n=3 independent biological replicates; Values are presented as the mean ± SEM; Two-Way ANOVA, p**<0.005; ***<0.0002; ****<0.0001; ns, not significant). **J.** Top: Confocal images of GFP- and GFP-(GR)_50_-transfected HEK-293 cells immunolabeled with an anti-rRNA antibody. Hoechst 33342 was used to counterstain nuclei. Scale bar = 20μm. Right: Dot plots showing the fluorescence intensity of rRNA staining in the whole cell (left) or the nuclear/cytoplasmic ratio (right) in GFP- and GFP-(GR)_50_-transfected HEK-293 cells. Individual cells are represented as dots from 3 independent experiments; bars represent median; Mann-Whitney U test, p****<0.0001).

**Supplementary Figure 3.**
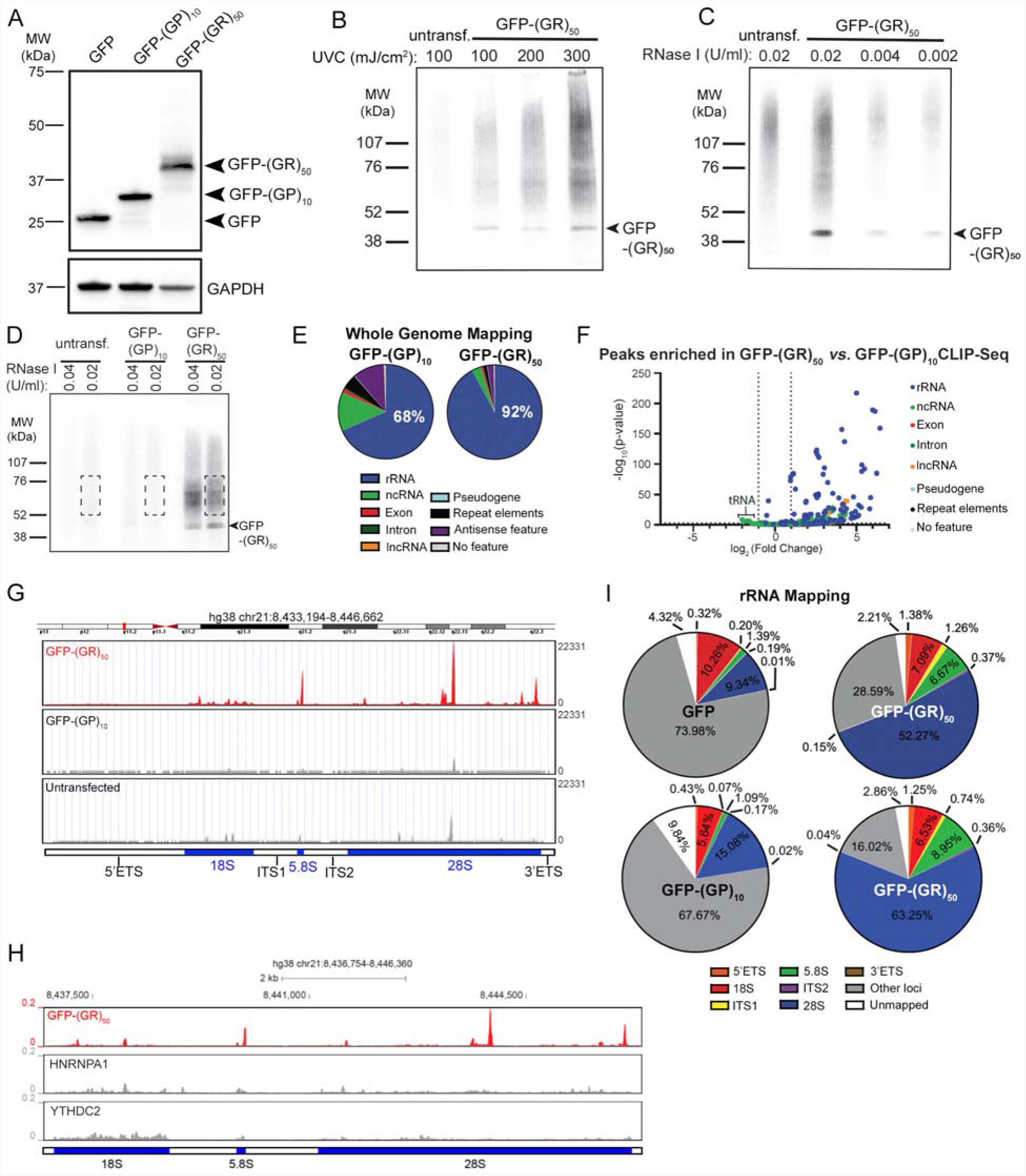
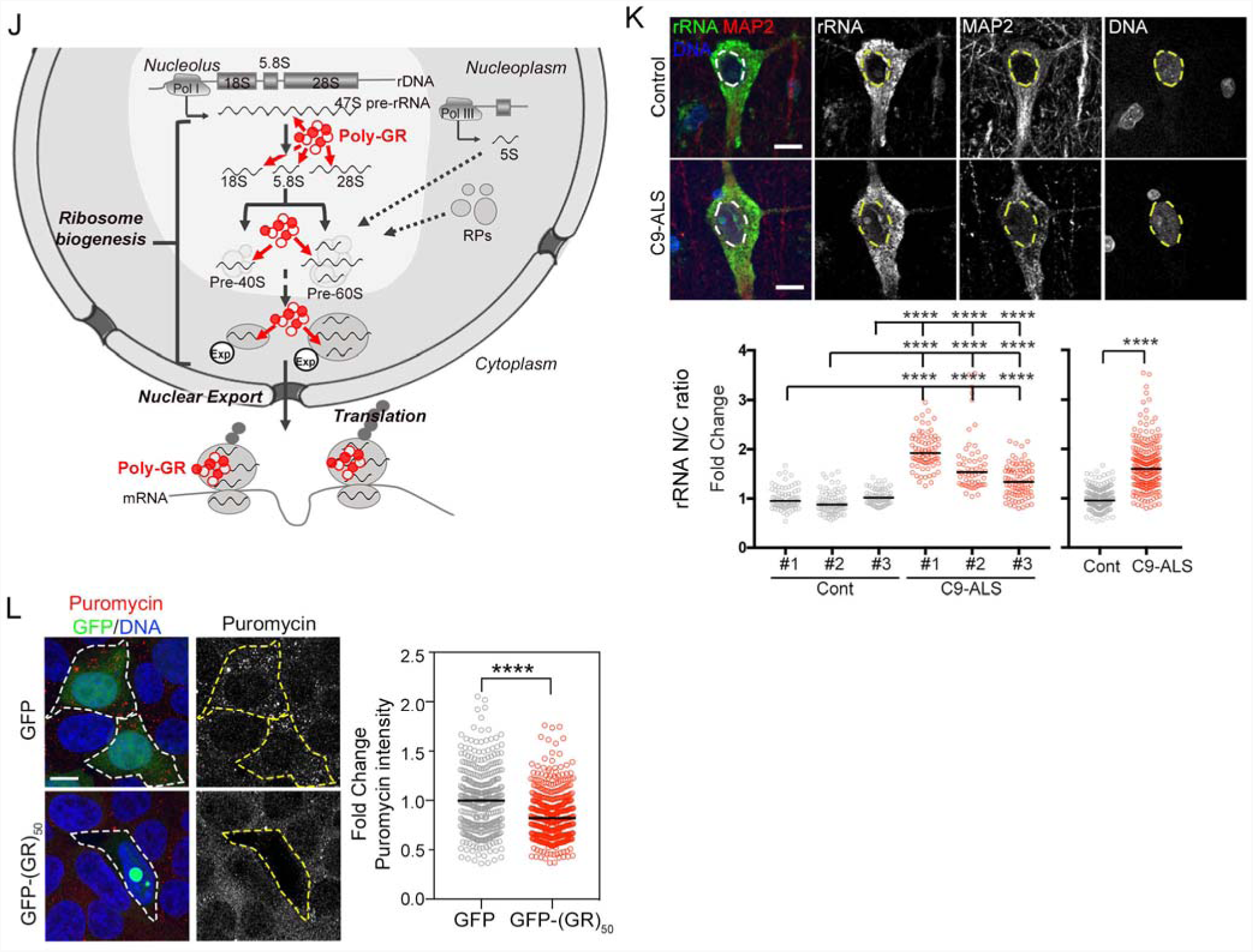
Characterization of the interactions between poly-GR and RNA in live cells. **A**. Lysates of HEK-293 cells transfected with GFP, GFP-(GP)_10_ or GFP-(GR)_50_ were subjected to WB to detect GFP. GAPDH is a loading control. **B**. After crosslinking HEK-293 cells expressing GFP-(GR)_50_ with increasing doses of UV (254 nm) and immunoprecipitation with anti-GFP antibodies, RNPs were labeled with [γ-^32^P]-ATP, fractionated by SDS-PAGE, transferred to a membrane and detected by autoradiography. Untransfected cells were controls. Arrow indicates the GFP-(GR)_50_ band. **C**. Lysates of UV-crosslinked untransfected HEK-293 cells and cells expressing GFP-(GR)_50_ were incubated with the indicated amounts of RNase I prior to immunoprecipitation and analysis as in (**B**). Arrow indicates the GFP-(GR)_50_ band. **D**. After UV crosslinking, HEK-293 cells expressing GFP, GFP-(GP)_10_ or GFP-(GR)_50_ were lysed, treated with the indicated amounts of RNase I and immunoprecipitated with anti-GFP antibodies. RNPs in immunoprecipitants were labeled with [γ-^32^P]-ATP and detected as in (**B)**. Dashed lines indicate RNPs excised for library preparation. Arrow indicates the GFP-(GR)_50_ band. **E**. Pie charts indicating the percentage of CLIP-Seq (replicate #2) peaks in the indicated classes of RNAs in GFP-(GP)_10_ and GFP-(GR)_50_ samples. Peaks were defined as having at least 5 overlapping reads. **F**. Volcano plot showing CLIP-Seq peaks enriched in GFP-(GR)_50_ compared to GFP-(GP)_10_ in CLIP-Seq (replicate #2). Negative log_2_ (fold change) indicates peaks that were more abundant in the GFP-(GP)_10_ control. As in replicate #1, GFP-(GR)_50_-CLIP-Seq peaks are enriched for rRNA, while tRNA peaks are more abundant in the control. **G**. Distribution of reads mapping to the RNA45SN1 locus from the GFP-(GR)_50_ (top), GFP-(GP)_10_ (middle) and untransfected (bottom) CLIP-Seq (replicate #2) analyses. **H**. Distribution of reads mapping to the RNA45SN1 locus from CLIP-Seq analyses after immunoprecipitation for GFP-(GR)_50_ (top), HNRNPA1 (middle) and YTHDC2 (bottom). **I**. Reads from the GFP, GFP-(GP)_10_ and GFP-(GR)_50_ CLIP-Seq (replicates #1 and #2) analyses were mapped to the RNA45SN1 locus. Reads that did not map to this locus were re-mapped to the hg38 human genome to determine the fraction that mapped to other genomic loci. Reads that did not map to hg38 are designated as “unmapped”. The percentage of reads in each category was quantified as a percentage of the total reads. **J**. Schematic model summarizing poly-GR interactions with rRNA and the potential consequences, including defects in ribosome biogenesis and protein translation. **K**. Top: Representative immunohistochemistry confocal images of layer V neurons immunolabeled for rRNA (green), MAP2 (red), and DNA (blue) in motor cortex tissue from a non-neurological control and a C9-ALS patient. Scale bars= 10μm. Bottom-left: Dot plot displaying the fold change in the N/C ratio of rRNA signal observed in cortical neurons of three non-neurological age-matched controls and three C9-ALS patients. Compiled control vs. C9-ALS data is shown on the bottom-right dot plot. Bars represent median; Mann-Whitney U test, p****<0.0001. **L**. Left: Confocal images of GFP- and GFP-(GR)_50_-transfected HEK-293 cells immunolabeled for puromycin. Hoechst 33342 was used to counterstain nuclei. Scale bar= 20μm. Right: Dot plot showing the fluorescence intensity of puromycin staining in GFP- and GFP-(GR)_50_-transfected HEK-293 cells. Individual cells are represented as dots from 3 independent experiments; bars represent median; Mann-Whitney U test, p****<0.0001.

Human rDNA exists as hundreds of copies of tandem repeats on five chromosomes (Potapova and Gerton, 2019). Although these repeats are not included in the assembled hg38 genome, five complete rDNA sequences that exist outside these clusters are included (RNA45SN1-5). Additionally, the genome is littered with divergent and truncated rRNA sequences that do not code for *bona fide* rRNA (Lander et al., 2001). Thus, to simplify quantification of the rRNA-derived reads, we remapped data from the CLIP experiments to a single full-length rDNA sequence, the RNA45SN1 locus (NCBI Accession NR 146117.1). We found that 69% and 81% of the total reads aligned to rDNA for the GFP-(GR)_50_ libraries (Figure S3H). In contrast, only 22% of the reads in the GFP control and 23% of the GFP-(GP)_10_ control libraries mapped to rDNA (Figure S3H). Interestingly, the overwhelming majority of GFP-(GR)_50_ reads were derived from the 28S region of the rDNA, with 52% and 63% of the total reads in the two libraries aligning to this region. Within the 28S rRNA, a prominent peak was located between nucleotides 2744 and 2810. This 66-nucleotide region was the most highly enriched site in both CLIP experiments accounting for 30% and 45% of total reads in the GFP-(GR)_50_ samples. An additional 57-nucleotide peak at the 3’ end of the 5.8S rRNA accounted for approximately 7% and 9% of the reads in the two GFP-(GR)_50_ datasets. Since mapping to rRNA is often considered nonspecific in CLIP-Seq experiments due to the abundance of rRNA, as an additional control we compared our results with rRNA-derived reads in datasets that were generated using the same iCLIP protocol (Huppertz et al., 2014). We asked whether iCLIP datasets examining two abundant RNA-binding proteins, HNRNPA1 or YTHDC2, contain similar rRNA-derived reads as poly-GR. We did not observe any substantial overlap in the sites of interaction, further suggesting that the binding between rRNA and GFP-(GR)_50_ we identified is highly specific (Figure S3I).

Since the production of mature rRNA involves a complex series of processing steps that occur almost entirely within nuclei, the GFP-(GR)_50_ CLIP peaks that we detected mapping to the 28S and 5.8S sites in the rDNA could be derived from nuclear precursors such as 47S, 45S, 30S, or 32S, and/or the mature cytoplasmic forms (5.8S, 28S) (Figure 3F). To further characterize the bound rRNAs and distinguish precursors from mature rRNA species, we used ribonucleoprotein immunoprecipitation (RIP) followed by northern blot (NB) analysis. These experiments revealed that mature 28S, 18S and 5.8S rRNAs were all detected in GFP-(GR)_50_, but not in GFP immunoprecipitates, as would be expected if mature cytoplasmic ribosomes were targets (Figure 3G-H). Consistent with specific binding, two other abundant RNAs, the SRP 7SL RNA and the RNase P RPPH1 RNA, were not detected in these immunoprecipitates. While these data demonstrate an association between GFP-(GR)_50_ and mature ribosomes, some CLIP reads could only have originated from immature rRNA (Figure S3H). The 5’ ETS and ITS1 regions were the most abundant precursor regions detected, each accounting for 0.7 to 1.3% of the reads in the GFP-(GR)_50_ CLIP datasets. Because these pre-rRNAs were not abundant enough to detect by NB, we assayed for their presence in our RIP experiment using RT-qPCR. Consistent with our NB results, RT-qPCR showed strong enrichment of mature 28S and 18S rRNAs in the GFP(GR)_50_ immunoprecipitants. This assay also demonstrated enrichment of sequences from the 5’ ETS and ITS1 regions of the pre-rRNA (Figure 3I). Noncoding Y RNAs and tRNA-Glu were selected as controls since these RNAs were not enriched in the GFP-(GR)_50_ CLIP-Seq dataset (Figure 3D). As expected, these RNAs showed no significant enrichment in the GFP-(GR)_50_ RIP (Figure 3I).

Collectively, these results demonstrate that poly-GR binds to multiple rRNA species, including precursors found in the nucleolus and mature RNAs found in both the nucleolus and within fully assembled cytoplasmic ribosomes (Figure S3J). These findings are well-aligned with the subcellular localization of GR in this *in vitro* cell model, a proportion of which accumulates within both the nucleolus and cytoplasm (Figure S2D). These interactions could potentially impact ribosomal assembly, homeostasis and/or function. In line with previous studies (Lee et al., 2016; White et al., 2019), immunolabeling to detect rRNA revealed reduced levels of total rRNA in cells transfected with poly-GR (Figure 3J). Importantly, we found a significant shift in the nucleocytoplasmic (N/C) ratio of rRNA signal in cells transfected with GFP-(GR)_50_ (Figure 3J). We also validated that this shift in rRNA occurs within cortical neurons in postmortem *C9ORF72* ALS/FTD tissue using immunohistochemistry (IHC) (Figure S4B). This increased N/C ratio of rRNA, which suggests impaired ribosomal homeostasis, likely contributes to the previously described reduced level of translation in poly-GR-containing cells (Figure S3L) (Hartmann et al., 2018; Lee et al., 2016; Loveland et al., 2022; Zhang et al., 2018b).

### The interaction of poly-GR with rRNA strongly contributes to ribosomal binding

The identification of rRNA as a binding partner for poly-GR is well aligned with previous mass spectrometry (MS)-based studies in cell models that have consistently uncovered ribosomal proteins (RPs) as GR-DPR interactors after immunoprecipitation (Figures 4A and S4A; Table S3) (Hartmann et al., 2018; Lee et al., 2016; Lopez-Gonzalez et al., 2016; Radwan et al., 2020; Tao et al., 2015; Yin et al., 2017; Zhang et al., 2018b). Similarly, IHC analyses in animal models and postmortem patient tissue, have shown strong co-localization between GR and RPs in neurons (Choi et al., 2019; Hartmann et al., 2018; Zhang et al., 2018b). In fact, 89% and 94% of all RPs of the small and large ribosomal subunits respectively, have been reported to precipitate along with poly-GR in at least one of the previously reported interactomes (Figures S4A-B). Intriguingly, one study showed that more than 40% of interactions with RP subunits disappear after RNA degradation by RNase treatment, suggesting that rRNA might be playing a critical role in the physical association of the ribosome with C9 poly-GR DPR (Figure S4C) (Lopez-Gonzalez et al., 2016). To investigate the significance of the GR-rRNA interaction more directly, we sought to model the poly-GR-ribosome complex. We used cryo-EM-based structural information of the mammalian ribosome (PDB ID: 5LZS) (Shao et al., 2016), and observed that some regions of the 5.8S and 28S rRNA, which were crosslinked to poly-GR in live cells, are exposed to the surface of the fully-assembled ribosome (Figures 4A-B). Interestingly, one of the closest RPs within this region is RPL7A/eL8, which is the only subunit that has been identified in all seven published poly-GR interactome experiments, and one that we validated in our model system (Figures 4A and S4A, D-E) (Hartmann et al., 2018; Lee et al., 2016; Lopez-Gonzalez et al., 2016; Radwan et al., 2020; Tao et al., 2015; Yin et al., 2017).

**Figure 4.**
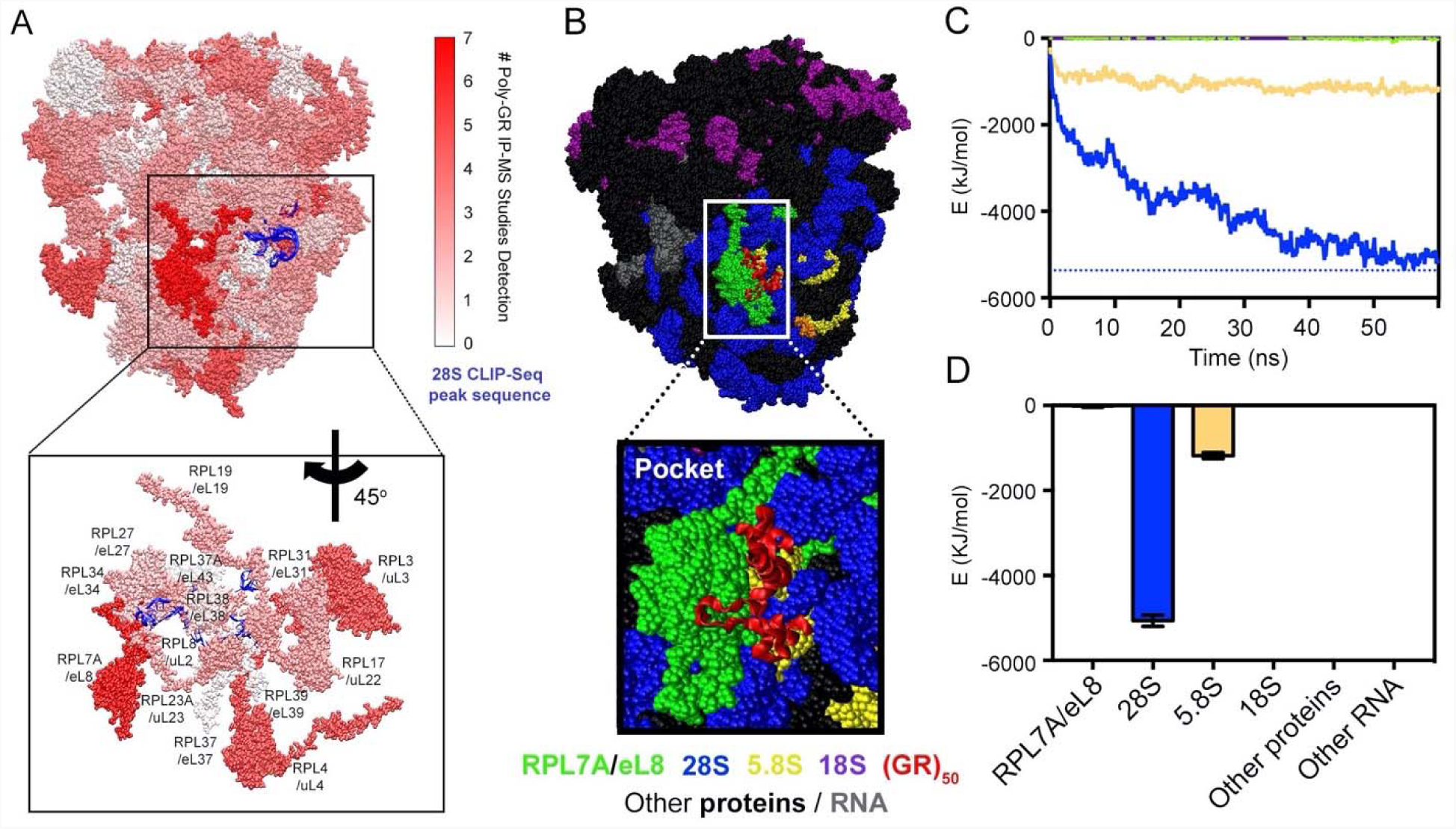
Computational modeling of poly-GR binding to ribosomal RNA and protein subunits. **A**. Top: Representative image recreated by all-atoms molecular dynamics simulations of all mammalian ribosomal proteins colored based on the number of studies that identified them as poly-GR interactors. The 28S rRNA sequence identified by our CLIP-Seq analysis is displayed in blue. Bottom: higher magnification of the mammalian ribosome proteome showing a region around the only protein that was co-precipitated with poly-GR in all 7 studies, RPL7A/eL8, which is in close proximity to the 28S rRNA sequence identified by our CLIP-Seq analysis. **B**. Representative image recreated by all-atoms molecular dynamics simulations of (GR)_50_ interacting with the mammalian ribosome. Based on our CLIP-Seq data and previous proteomic data (IP-MS), we determined a poly-GR interaction region (“pocket”) magnified in the inset at the bottom. The targeted RPs and RNAs were color labeled as follows: RPL7A:green; RPS30/RPL37/RPL39/RPL41:orange; 28S:blue; 5.8S:yellow; 18S:purple; Other proteins and RNAs are labeled in black and gray respectively. **C**. Line bars indicating the energetic strength of the interactions of (GR)_50_ in the “pocket” region of the ribosome over time. E>0 indicates repulsion; E<0 indicates attraction. The energy of interaction for 28S (blue), 5.8S (yellow), 18S (purple) and RPL7A (green) is shown by each line. Dashed line indicates the highest interaction energy achieved between (GR)_50_ and a 28S rRNA. **D**. Bar graphs showing average energy of interaction of (GR)_50_ with different ribosomal components as in **C** during the last 10 ns, when structure is considered stable. Values are presented as mean ± SD.

**Supplementary Figure 4.**
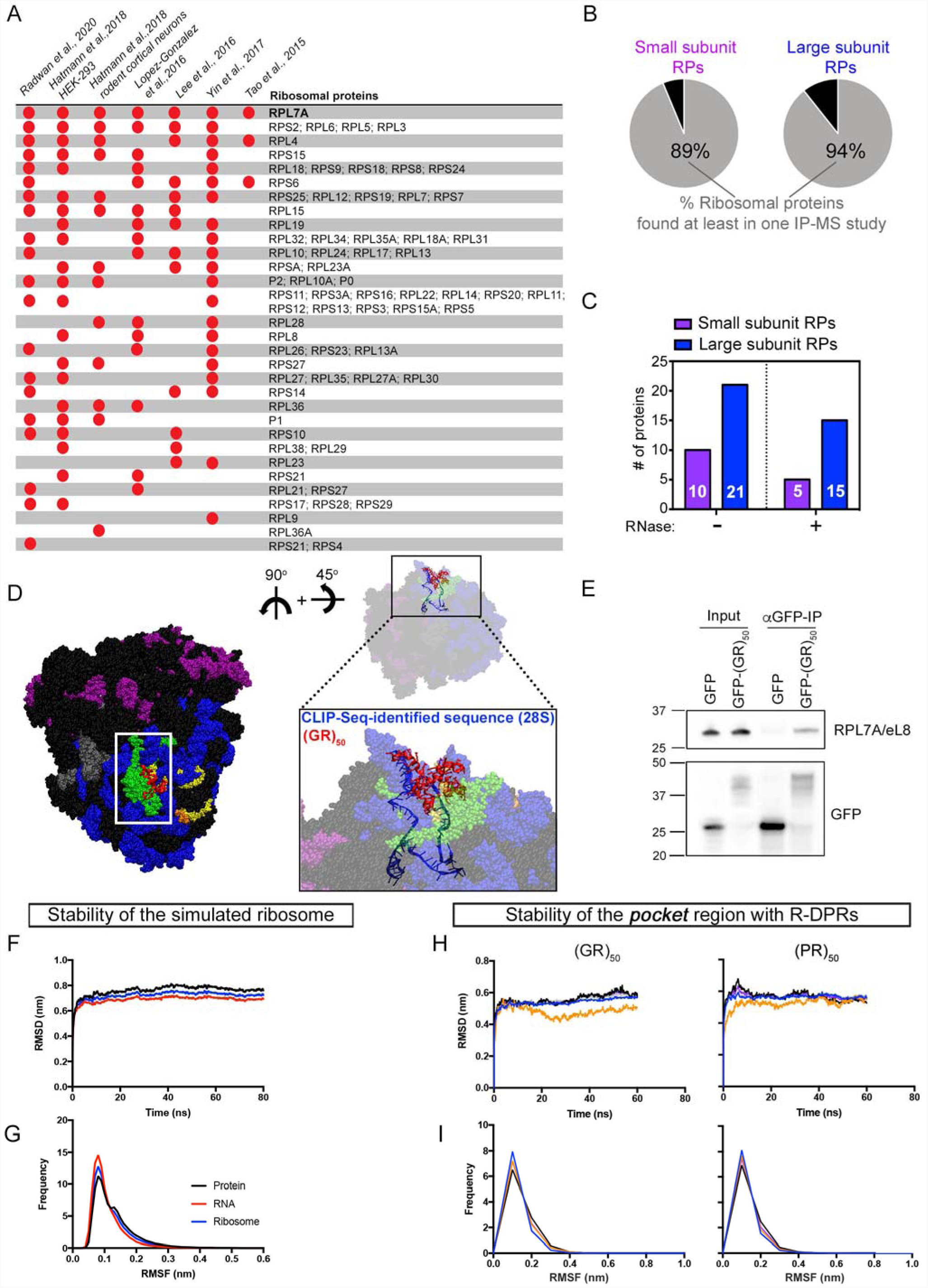
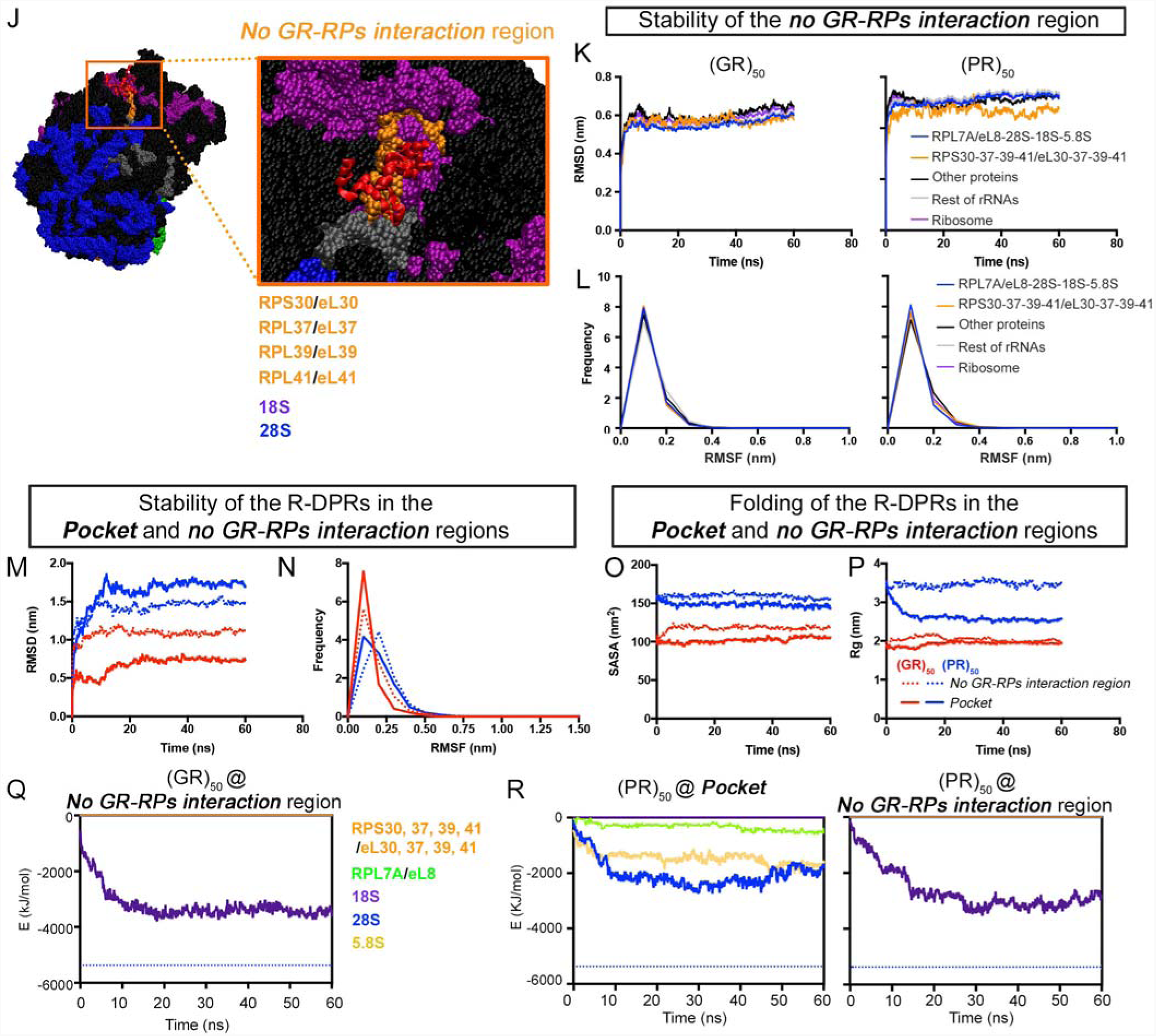
Computational modeling of poly-GR binding to ribosomal RNA and protein subunits. **A**. Comparative data analysis of the ribosomal proteins (RPs) detected in published datasets of IP-MS experiments performed in mammalian *in vitro* cell models. Red dots indicate a positive interaction of poly-GR with the respective RPs. **B**. Pie charts showing the percentage of small and large ribosomal subunit proteins immunoprecipitated with poly-GR in at least one previous study. **C**. Bar graphs displaying the number of small (purple) and large (blue) ribosomal subunit proteins immunoprecipitated with poly-GR in the presence or absence of RNase as described by (Lopez-Gonzalez et al., 2016). **D**. Left: All-atoms molecular dynamics simulation of the mammalian 80S ribosome showing a putative region (indicated by a white line box) of interaction with (GR)_50_ (pocket). The RPs and RNAs surrounding this region are labeled as follows: RPL7A: green; RPS30/RPL37/RPL39/RPL41: orange; 28S: blue; 5.8S: yellow; 18S: purple; Other proteins and RNAs are labeled in black and gray respectively. (GR)_50_ is colored in red. Right: A different angle of view of the mammalian ribosome reflects the close proximity between the 28S sequence identified by our CLIP-Seq analysis and RPL7A/eL8, consistently detected as a (GR)_50_-interactor by 7 independent IP-MS studies. Angle of view with respect to the image on the left is represented at the top right part of the panel **E**. Immunoprecipitation (IP)-WB analysis of GFP-(GR)_50_ or GFP in HEK-293 transfected cells. Cell lysates were subjected to IP with anti-GFP antibodies and WB was performed to detect RPL7A/eL8 (top) and GFP (bottom). **F**. Line graphs showing changes from the initial structure calculated by root-mean-square deviation (RMSD) over time in the different components of the mammalian ribosome. **G**. Line graphs showing the frequency of changes calculated by root-mean-square fluctuation (RMSF) of the last 10 ns in the different components of the mammalian ribosomal pocket region. **H**. Line graphs showing changes from the initial structure calculated by RMSD over time in the RNA and protein components of the “*pocket”* region of the mammalian ribosome in the presence of (GR)_50_ and (PR)_50_. **I**. Line graphs showing the frequency calculated by RMSF of the last 10 ns in the different components of the mammalian ribosomal “*pocket”* region in the presence of (GR)_50_ and (PR)_50_. **J**. Representative image recreated by all-atoms molecular dynamics simulations of (GR)_50_ and (PR)_50_ interacting with distinct rRNAs and RPs present in the “*no GR-RPs interaction”* region magnified in the inset on the right. Based on 7 different IP-MS studies, the proteins in this ribosomal region do not interact with poly-GR. **K**. Line graphs showing changes from the initial structure calculated by RMSD over time in the RNA and protein components of the “*no GR-RPs interaction”* region of the mammalian ribosome in the presence of (GR)_50_ and (PR)_50_. **L**. Line graphs showing the frequency of changes of the last 10 ns in the RNA and protein components of the “*no GR-RPs interaction”* region of the mammalian ribosome in the presence of (GR)_50_ and (PR)_50_. **M**. Line graphs showing changes from the initial structure calculated by RMSD over time of (GR)_50_ and (PR)_50_ in the “*pocket”* and “*no GR-RPs interaction”* region of the mammalian ribosome. **N**. Line graphs showing the frequency of changes calculated by RMSF of the last 10 ns for (GR)_50_ and (PR)_50_ in the “*pocket”* and “*no GR-RPs interaction”* region of the mammalian ribosome. **O**. Line graphs representing the SASA value changes over time of (GR)_50_ and (PR)_50_ in the “*pocket”* and “*no GR-RPs interaction”* region of the mammalian ribosome. **P**. Line graphs representing the radius gyration changes over time of (GR)_50_ and (PR)_50_ in the “*pocket”* and “*no GR-RPs interaction”* region of the mammalian ribosome. **Q**. Line bars indicating the energetic strength of the interactions of (GR)_50_ in the “*no GR-RPs interaction”* region of the ribosome over time. Dashed line indicates the highest interaction energy achieved between (GR)_50_ and 28S rRNA. **R**. Line bars indicating the energetic strength of the interactions of (PR)_50_ within the “pocket” (left) and the “*no GR-RPs interaction”* (right) region of the ribosome over time. Dashed line indicates the highest interaction energy achieved between (GR)_50_ and 28S rRNA. Raw data of all the simulations presented in this figure can be found in Table S4.

Based on this converging evidence from proteomic and transcriptomic experiments, we established an *in-silico* model and mapped one of likely many regions that poly-GR can bind to, on the surface of the ribosome (Figures 4B and S4F-I). This potential binding “pocket” within the modelled poly-GR-ribosome assembly, accommodates the physicochemical interaction of poly-GR with both rRNA and RPs. To better understand the dynamics of this interaction we performed molecular dynamics (MD) simulations. We specifically asked how strongly the different ribosomal components that are aligned either within the pocket (RPL7A/eL8, 5.8S, 28S and 18S), or in close proximity to the pocket (other proteins and RNA), contribute to the binding with (GR)_50_ by comparing their energy of interaction (E<0 attractive and E>0 repulsive) (Figures 4B-D and S4F-I, M-P). Intriguingly, this analysis showed that the 28S rRNA had the highest energy of interaction with (GR)_50_ (E = −4004.79 kJ/mol), followed by 5.8S (E = −1059.01 KJ/mol), while RPL7A/eL8 and 18S exhibited negligent E values (Figures 4C-D). We also simulated the interaction of (GR)_50_ in a different ribosomal region, where there are RPs that have never been immunoprecipitated with poly-GR (no GR-RPs interaction control region) (Figures S4J-K, M-P). We found that (GR)_50_ interacted exclusively with 18S rRNA in this region, although with weaker energy of interaction relative to the one exhibited for 28S rRNA in the pocket region (Figures 4C and S4Q). Of note, (GR)_50_ acquires a more folded configuration within the pocket than within the control region (Figures S4O-P), suggesting a higher level of conformational adaptation in the pocket, likely due to its stronger interaction with the 28S rRNA. Thus, our computational and CLIP-Seq data (Figures 3 and 4) suggest that interactions between poly-GR and rRNA contribute to ribosomal binding.

### Custom rRNA oligonucleotides bind poly-GR and inhibit the toxic effects of poly-GR in multiple model systems

Collectively, our data indicate that the physicochemical properties of poly-GR promote strong interactions with RNA, while independent transcriptomic and proteomic analysis suggest that GR-DPR can interact with multiple ribosomal RNA species and protein subunits (Figures 3–4 and S3–S4). Importantly, these interactions can take place with both assembling ribosomal subunits in the nucleus and mature cytoplasmic ribosomes (Hartmann et al., 2018; Zhang et al., 2018b), and likely contribute to an impairment in ribosomal homeostasis and function (Figure 3J and S4). Our dynamic simulation models suggest that the strong rRNA-GR binding may mediate the interaction between the R-DPR and the ribosome. We thus reasoned that an RNA molecule of the right sequence and structure could act as a “bait” for poly-GR, binding to it and sequestering it away from pathological interactions with other proteins and RNAs.

To test this hypothesis, we used the 28S rRNA sequence that was significantly enriched in our CLIP-Seq experiments and designed an oligonucleotide RNA bait with 2’-O-methyl modifications in both the 5’ and 3’ ends to enhance its stability and binding properties (Figure 5A and Methods). We first confirmed that the bait could interact with synthetic (GR)_15_ *in vitro* (Figure S5A), as well as with GFP-(GR)_50_ in live transfected mammalian cells (Figures 5B-C). Immunoprecipitation of GFP or GFP-(GR)_50_ from whole cell extracts coupled to NB analysis, demonstrated a specific interaction between the 28S rRNA-based bait and GFP-(GR)_50_ but not with the GFP control (Figure 5B). To quantitate the specificity of this interaction in a different way, we co-transfected GFP or GFP-(GR)_50_ cells with the 28S rRNA-Cy3 bait or a scrambled RNA-Cy3 molecule of the same length and percentage of nucleotide composition. Quantitative measurements of Cy3 fluorescence after precipitation with a GFP antibody demonstrated a specific interaction between GFP-(GR)_50_ and the 28S bait but not with the scrambled control (Figure 5C).

**Figure 5.**
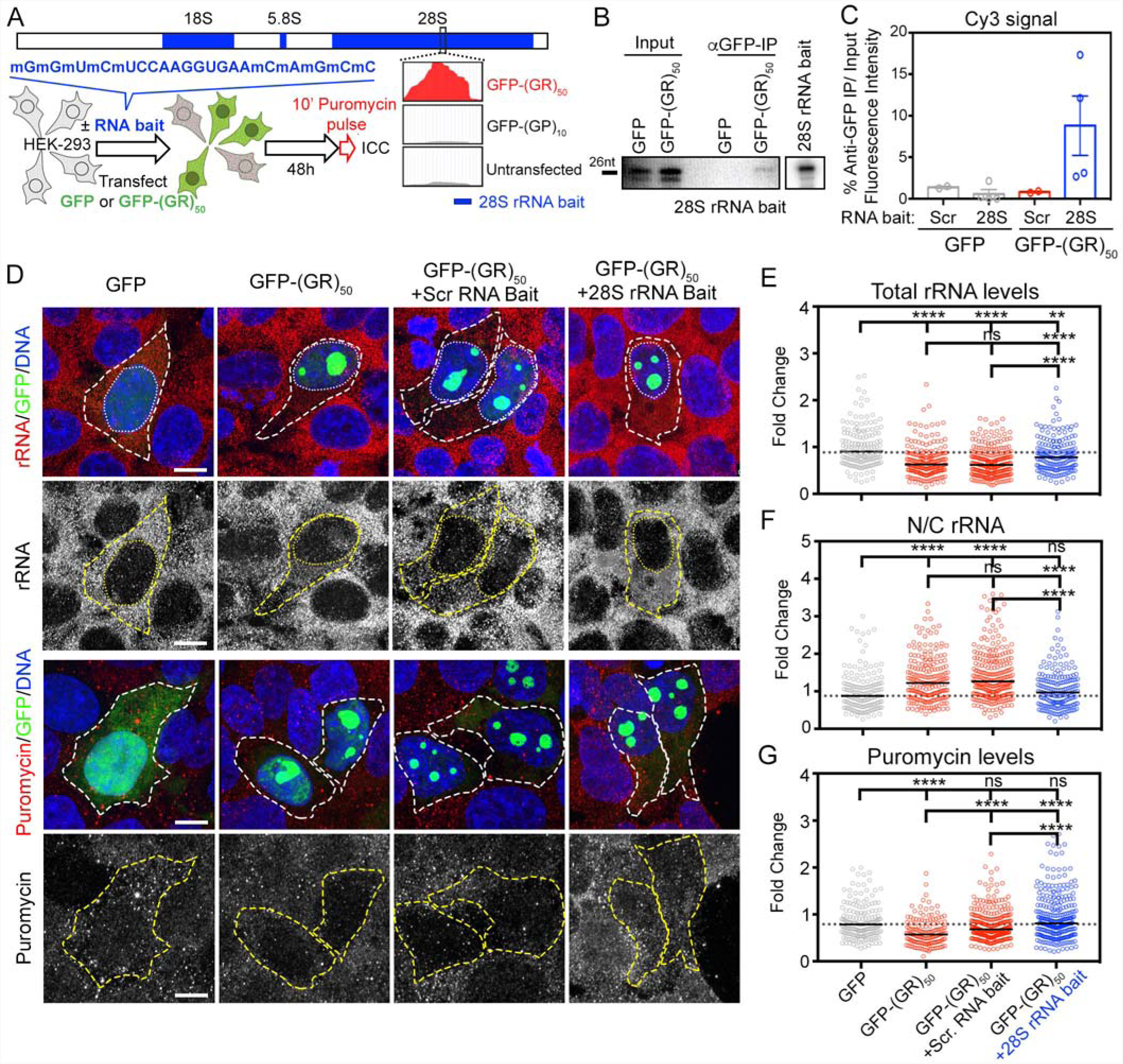
Modified RNA-based strategy to inhibit the toxic effects of poly-GR on ribosomal homeostasis. **A**. Left: Schematic showing the modified RNA blocking strategy based on the findings of the CLIP-Seq analysis. Right: Schematic representing the experimental workflow for rescue experiments utilizing the modified 28S rRNA baits. **B**. Lysates of HEK-293 cells co-transfected with 28S rRNA-based bait and GFP-(GR)_50_ or GFP were subjected to immunoprecipitation with an anti-GFP antibody. RNAs within immunoprecipitants were fractionated in denaturing polyacrylamide/urea gel and the indicated 28S rRNA bait was detected by NB exclusively in the GFP-(GR)_50_ condition. The input represents 1% of the lysate used in immunoprecipitation. **C**. Lysates of HEK-293 cells co-transfected with 28S rRNA or scrambled control bait and GFP-(GR)_50_ or GFP were subjected to immunoprecipitation with an anti-GFP antibody. Bar graphs showing the level of Cy3-labeled bait detected within the immunoprecipitants normalized to input. **D**. Confocal images of GFP- and GFP-(GR)_50_-transfected HEK-293 cells immunolabeled for rRNA (top) and puromycin (bottom). Hoechst 33342 was used to counterstain nuclei. Scale bar= 10μm. **E**. Dot plots showing the fluorescence intensity of rRNA staining in GFP- and GFP-(GR)_50_-transfected HEK-293 cells treated with scrambled or 28S rRNA-based RNA baits. **F**. Dot plots showing the N/C rRNA ratio in GFP- and GFP-(GR)_50_-transfected HEK-293 cells treated with scrambled or 28S rRNA-based baits. **G**. Dot plots showing the fluorescence intensity of puromycin staining in GFP- and GFP-(GR)_50_-transfected HEK-293 cells treated with scrambled or 28S rRNA-based baits. In D-F, individual cells from 3 independent experiments are represented as dots, the dotted line marks the median in GFP-transfected cells, and bars represent the median; Mann-Whitney U test, p**<0.01; ****<0.0001; ns, not significant).

**Supplementary Figure 5.**
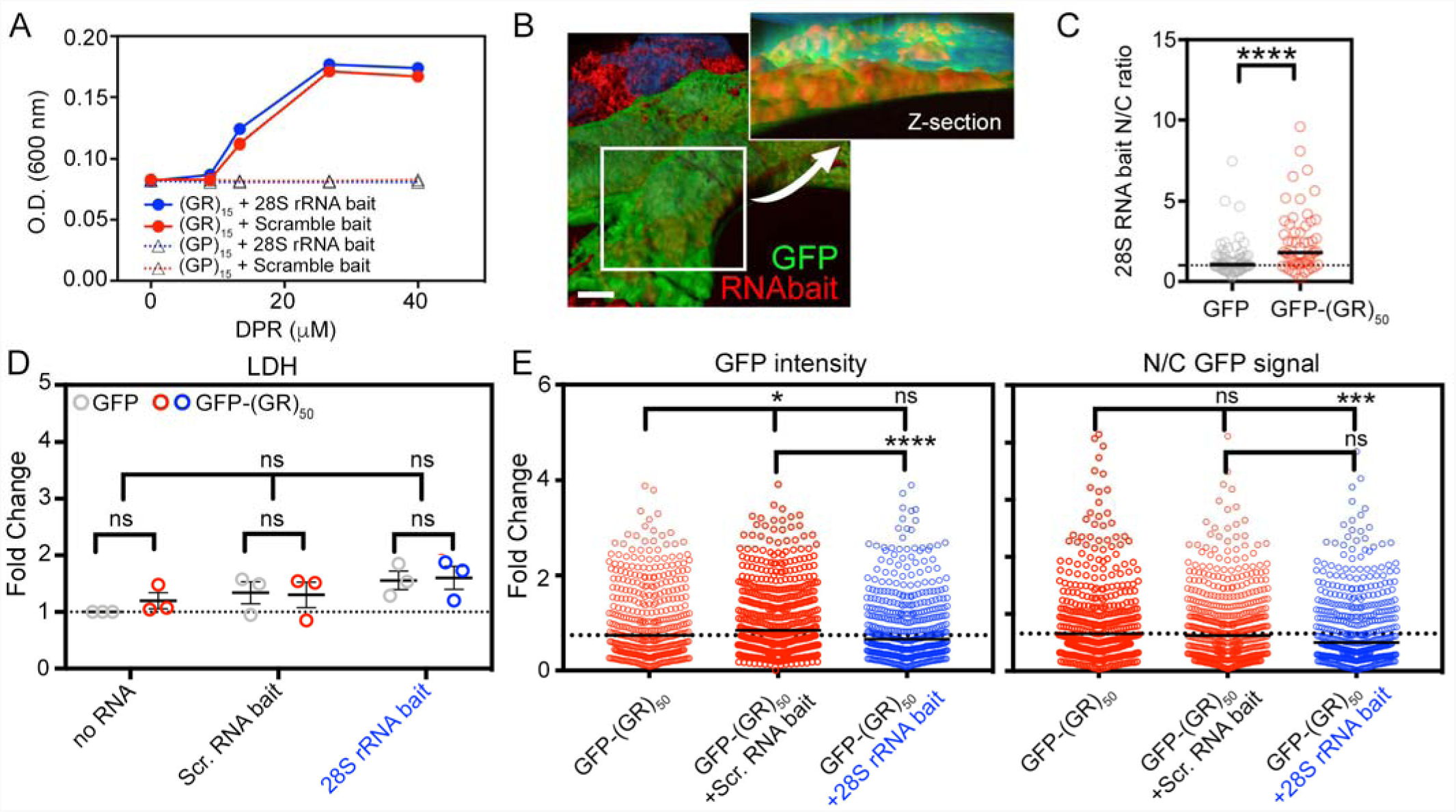
Modified RNA-based strategy to inhibit the toxic effects of poly-GR on ribosomal homeostasis. **A**. Line graphs depicting DPR ((GR)_15_ and (GP)_15_) concentration-dependent precipitation of 28S and scramble rRNA-based baits calculated by optical density. **B**. Volumetric reconstruction of confocal images of HEK-293 cells sequentially co-transfected with 28S rRNA-based bait conjugated with Cy3 (red) and GFP (green). Hoechst 33342 was used to counterstain nuclei. Scale bar= 10μm. Inset shows a single plane reconstruction of GFP and Hoechst 33342 signal in combination with volumetric Cy3-bait signal. **C**. Dot plot showing the nuclear/cytosolic (N/C) distribution of the RNA bait conjugated with Cy3 in GFP- or GFP-(GR)_50_-transfected HEK-293 cells. Individual cells are represented as dots from 3 independent experiments and bars represent median. Mann-Whitney U test, p***<0.001; p****<0.0001. **D**. Dot plot showing the levels of LDH in the media of HEK-293 cells sequentially co-transfected with scrambled or 28S-rRNA based bait and GFP- or GFP-(GR)_50_. Average values of each experiment are represented as dots and the dotted line marks the median in no RNA bait treated GFP-transfected cells (n=3 independent biological replicates; bars represent mean ± SEM; ANOVA, ns, not significant). **E**. Dot plot showing the intensity (left) and N/C distribution (right) of GFP signal in GFP-(GR)_50_-transfected HEK-293 cells upon treatment with scrambled or 28S rRNA-based bait. Individual cells are represented as dots and the dotted line marks the median in no RNA bait-treated GFP-(GR)_50_-transfected cells (n=3 independent biological replicates; bars represent median; Mann-Whitney U test, p*<0.05; p***<0.001; p****<0.0001; ns, not significant).

Next, we delivered the 28S and scrambled control baits to cells expressing GFP or GFP-(GR)_50_ and monitored several metrics related to poly-GR and its adverse effects on ribosomal homeostasis. First, we used confocal imaging to establish that the baits were effectively internalized (Figure S5B) and observed that the 28S rRNA-Cy3 bait exhibited higher accumulation in the nucleus of cells expressing GFP-(GR)_50_ relative to cells expressing GFP, likely on account of the interaction with nuclear poly-GR (Figure S5C). Critically, we did not observe any adverse toxic effects in cells treated with either the 28S rRNA-based or scrambled-RNA bait (Figure S5D). The 28S rRNA bait resulted in a moderate but significant reduction in the N/C ratio of the GFP-GR signal relative to the untreated poly-GR overexpressing cells, suggesting that binding to the bait affected the localization of GR (n = 443 cells for no RNA bait; 572 cells for +Scr. bait; 491 cells for +28S rRNA bait; p < 0.001) (Figure S5E). As we described earlier, cells expressing GFP-(GR)_50_ exhibited a significant reduction in total levels of rRNA and increased rRNA N/C ratio (n = 157 cells for GFP; 213 cells for GFP-(GR)_50_; p < 0.0001), as well as a decrease in protein translation (n = 178 cells for GFP; 153 cells for GFP-(GR)_50_; p < 0.0001) (Figures 5D-G). Importantly, all these deficits were significantly inhibited by the presence of the 28S-based bait relative to untreated GFP-(GR)_50_ (n = 186 cells; p < 0.0001 for total rRNA levels and N/C rRNA ratio, and n = 306 cells; p < 0.0001 for protein translation), with rRNA localization and protein translation restored to levels that were similar to the ones seen in GFP-control cells (Figures 5D-G). This protective effect was specific to the 28S bait, as its beneficial effects were significantly different relative to cells treated with the scrambled control molecule (n = 269 cells; p < 0.001 for both total rRNA levels and N/C rRNA ratio; n = 303 cells; p < 0.0001 for protein translation) (Figures 5D-G).

We next interrogated the effects of the 28S bait on a neuronal model of poly-GR toxicity. We differentiated healthy control iPSCs into spinal motor neurons (MNs) using a well characterized protocol (Ziller et al., 2018), and transduced the cultures with a lentivirus expressing GFP or GFP-(GR)_50_. We used live cell imaging analysis to track individual cells over the course of 90 days and found that poly-GR overexpressing MNs exhibited significantly reduced survival relative to GFP-expressing MNs (n = 221 GFP MNs; and n = 223 GFP-(GR)_50_ MNs; 8-10% reduction, p = 0.0038) (Figure 6A). All degenerating neurons were characterized by the accumulation of poly-GR-GFP nuclear aggregates, and nuclear aggregation strongly predisposed MNs to degeneration (Figures 6B-C). Importantly, MNs with poly-GR-GFP nuclear aggregates progressively exhibited a significant reduction in total rRNA levels, accumulation of rRNA signal within the nucleus, and reduced protein translation (Figures 6D and S6A-B). Treating these MN cultures with the 28S rRNA bait led to a significant increase in survival (Figure 6E) that was associated with a 46% reduction in MNs with poly-GR nuclear aggregates (Figure 6F), and a 25% reduction in cell death within MNs with poly-GR nuclear aggregates (Figure 6G). These results suggest that the 28S bait mitigates toxicity by slowing down the accumulation of toxic nuclear poly-GR. In contrast, the scrambled control had only a minor effect on survival and did not rescue the accumulation of nuclear poly-GR-aggregates. To assess if the 28S bait can successfully mitigate the toxic effects of the *C9orf72* hexanucleotide repeat expansion (C9-HRE) in a more disease-relevant, physiological system, we used fibroblast-induced MNs (iMNs) (Shi et al., 2018), derived from C9-HRE ALS patients or healthy control subjects (Figures 6H, top). Strikingly, administration of single dose of 28S rRNA significantly improved survival and reduced the hazard ratio in iMNs derived from three distinct C9 ALS patients, while the scrambled control RNA had no substantial effect (Figures 6H-I).

**Figure 6.**
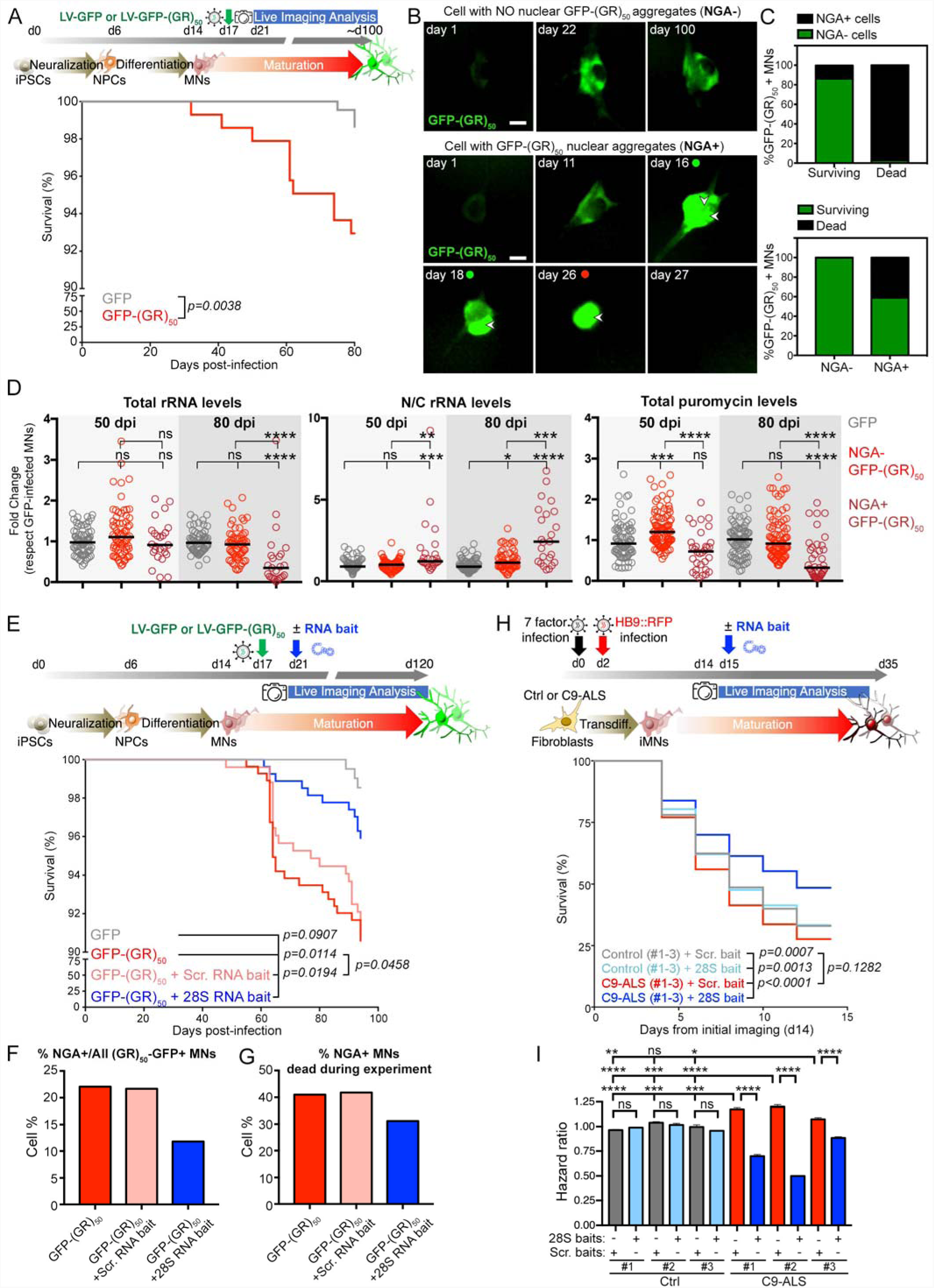
RNA-based strategy to inhibit the toxic effects of poly-GR in stem cell derived MNs. **A**. Top: Schematic representation of the experimental workflow for live imaging of iPSC-derived MNs transduced with GFP or GFP-(GR)_50_ lentiviruses. Bottom: Kaplan-Meier plot displaying survival of iPSC-derived MNs for >80 days after infection (dpi). Each trace includes neurons from three independent differentiations; n(GFP)=221 and n(GFP-(GR)_50_)=223 cells. Comparisons between conditions were assessed by Gehan-Breslow-Wilcoxon test; p-values are displayed in the graph. **B**. Images of GFP- and GFP-(GR)_50_-transduced MNs at different dpi. Arrowheads indicate the presence of nuclear GR aggregates (NGA). Green or red dots indicate the time at which NGAs appear and MN degenerate, respectively. Scale bar=10μm. **C**. Top: Bar plot showing the percentage of surviving and degenerating GFP-(GR)_50_-transduced MNs, with or without nuclear GR aggregates (NGA) and, bottom: the percentage of GFP-(GR)_50_-transduced MNs with or without NGAs that survive or degenerate. **D**. Dot plot showing the fold change in total rRNA, N/C rRNA and puromycin levels in GFP control and GFP-(GR)_50_-expressing MNs, with or without NGA, at 50 and 80 dpi. Each dot represents the value of a single MN. Comparisons between conditions were assessed by a Mann-Whitney U test, p*<0.05, p**<0.01, p***<0.001, p****<0.0001; ns, not significant. **E**. Top: Schematic representation of the experimental workflow for live imaging of iPSC-derived MNs transduced with GFP or GFP-(GR)_50_ lentiviruses treated with scramble- or 28S rRNA-based baits. Bottom: Kaplan-Meier plot displaying survival of iPSC-derived MNs for all four conditions. Each trace includes neurons from three independent differentiations; n(GFP)=206, n(GFP-(GR)_50_)=276, n(GFP-(GR)_50_ + Scramble bait)=277, n(GFP-(GR)_50_ + 28S rRNA-based bait)=269 cells. Comparisons between conditions were assessed by Gehan-Breslow-Wilcoxon test; p-values are displayed in the graph. **F**: Bar plot showing the percentage of MNs that exhibited nuclear GR aggregates (NGA+) over ~90 dpi. **G**. Bar plot showing the percentage of GFP-(GR)_50_ expressing MNs with or without NGA that degenerated over ~90 dpi. **H**. Top: Schematic representation of the experimental workflow for live imaging of induced MNs (iMNs) from three control and three C9orf72-ALS (C9-ALS)-patient derived iPSC lines, which were non-treated or treated with scramble- or 28S rRNA-based baits. Bottom: Kaplan-Meier plot displaying survival of iMNs for all four conditions. Each trace includes neurons from three independent iPSC lines; n=100 cells/cell line and condition. Comparisons between conditions were assessed by Gehan-Breslow-Wilcoxon test; p-values are displayed in the graph. **I**. Bar plot showing the hazard ratio of three control and three C9-ALS iMNs non-treated or treated with scramble- or 28S rRNA-based baits. Two sample comparisons between treated vs. non-treated conditions were performed by T-test, and multiple comparisons by ANOVA; p*<0.05; p**<0.01; ***<0.001; ****<0.0001; ns, not significant.

**Supplementary Figure 6.**
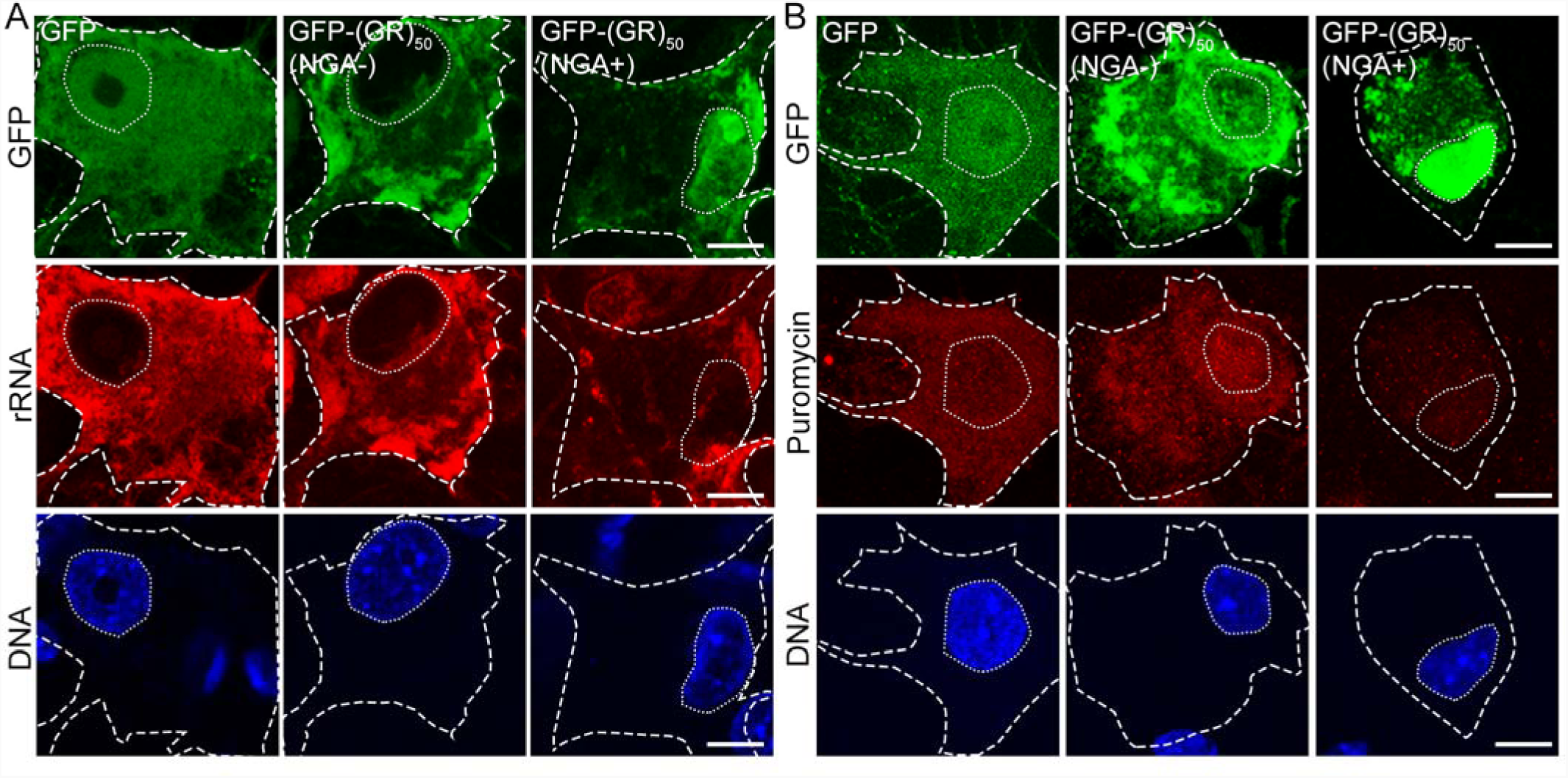
RNA-based strategy to inhibit the toxic effects of poly-GR in stem cell derived MNs. **A**. Confocal images of GFP- and GFP-(GR)_50_-transduced iPSC-derived MNs, with or without NGA, immunolabeled for rRNA at 80 days post-infection (dpi). Hoechst 33342 was used to counterstain nuclei. Scale bar = 10 μm. **B**. Confocal images of GFP- and GFP-(GR)_50_-transduced iPSC-derived MNs, with or without NGA, immunolabeled for puromycin at 80 days post-infection (dpi). Hoechst 33342 was used to counterstain nuclei. Scale bar = 10 μm.

Lastly, to assess the ability of the rRNA bait molecule to ameliorate poly-GR toxicity in an intact nervous system *in vivo*, we utilized two *Drosophila* models of poly-GR overexpression (Figures 7 and S7). Motor neuronal expression of (GR)_50_-EGFP is lethal during development, with ~99% of mutant flies dying at pupal stages and failing to eclose (n = 161 flies; Figure 7A-D). We reasoned that this highly toxic model would represent a stringent platform to test for any beneficial effects of the modified 28S RNA molecule on GR-DPR toxicity *in vivo*. We first ensured that the administration of Cy3-conjugated bait in feeding media during early larval stages, led to its sufficient uptake within larval brain cells (Figures 7A-B). Importantly, we found that the 28S rRNA-based bait had no effect on control GFP flies, while it significantly mitigated GR toxicity in (GR)_50_-EGFP mutant flies with up to 9.3% of animals effectively eclosing and reaching adult stages (Figures 7D) (n = 151-164 flies per group; p < 0.001). We obtained similar results using an alternative fly model, were (GR)_36_ is specifically expressed in the fly eye. Treatment with 28S rRNA-based bait, had a moderate but highly significant effect, reducing the severity of (GR)_36_-dependent eye degeneration and the appearance of necrotic patches (Figures S6A-C). Collectively, these experiments demonstrate that administration of a modified 28S rRNA bait molecule can significantly meliorate poly-GR-dependent toxicity in multiple model systems *in vitro* and *in vivo*.

**Figure 7.**
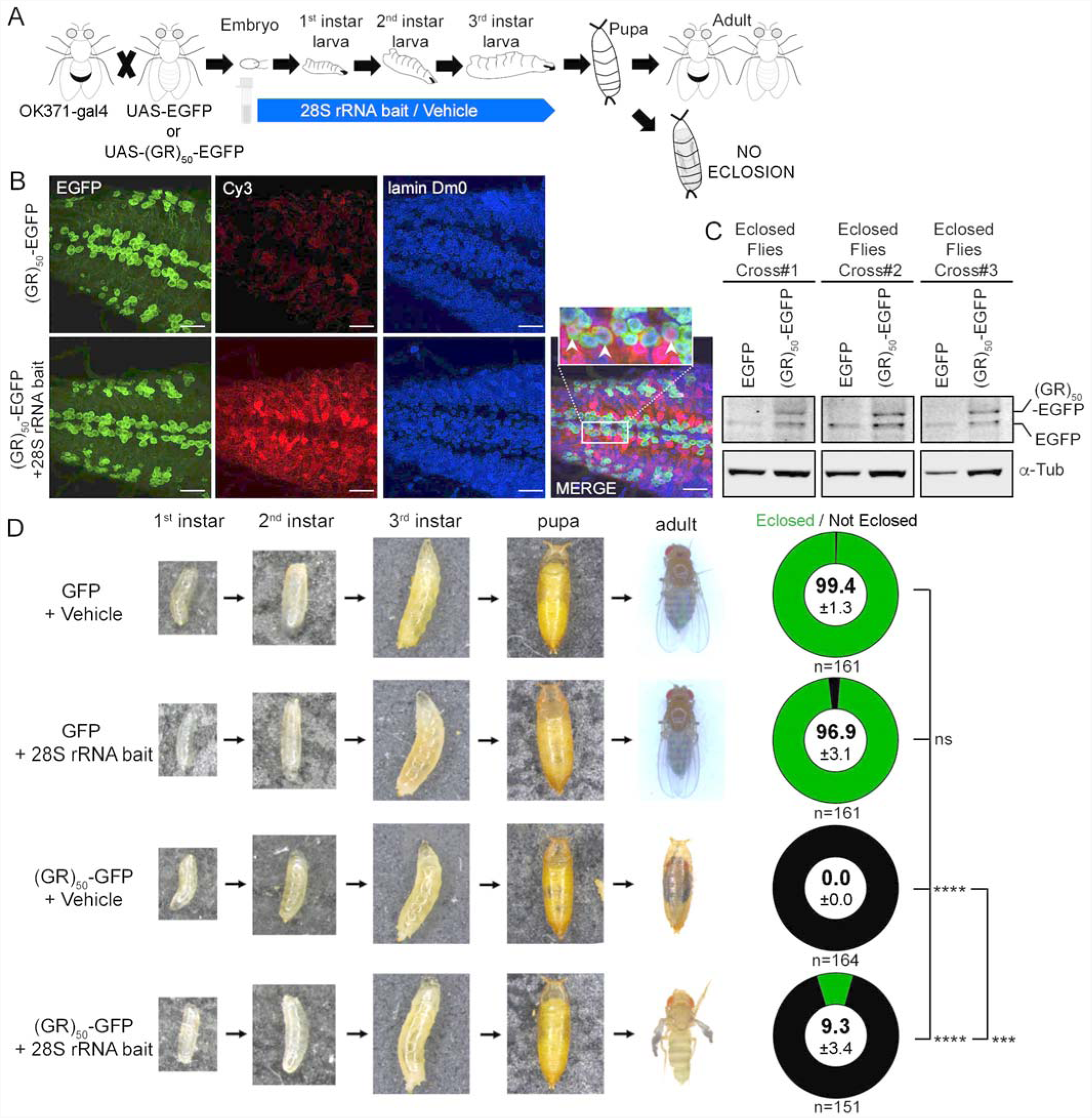
RNA-based strategy to inhibit the toxic effects of poly-GR *in vivo*. **A**. Schematic illustrating a *Drosophila* model to study poly-GR toxicity in the presence or absence of the 28S rRNA-based bait. **B**. Confocal images showing the GFP and (GR)_50_-GFP transgene expression and the presence of 28S rRNA-based bait conjugated with Cy3 in larval brains. Lamin Dm0 immunolabeling was used as a counterstaining. Arrowheads in merged image insert display the co-localization of (GR)_50_-GFP and 28S rRNA-based bait in larval brain cells. Scale bar= 10μm. **C**. Western blot depicting the protein expression of GFP and (GR)_50_-GFP transgenes in brains of flies treated with 28S rRNA-based bait that eclosed. Alpha-tubulin (α-tub) was used as a loading control. **D**. Representative images of larvae, pupa and adult (OK371-GAL4 x EGFP) or (OK371-GALl4 x (GR)_50_-EGFP) mutant flies in the presence or absence of the 28S rRNA-based bait. Pie charts on the right indicate the percentage of pupae eclosion in the different conditions. Number of animals analyzed per condition is also displayed.

**Supplementary Figure 7.**
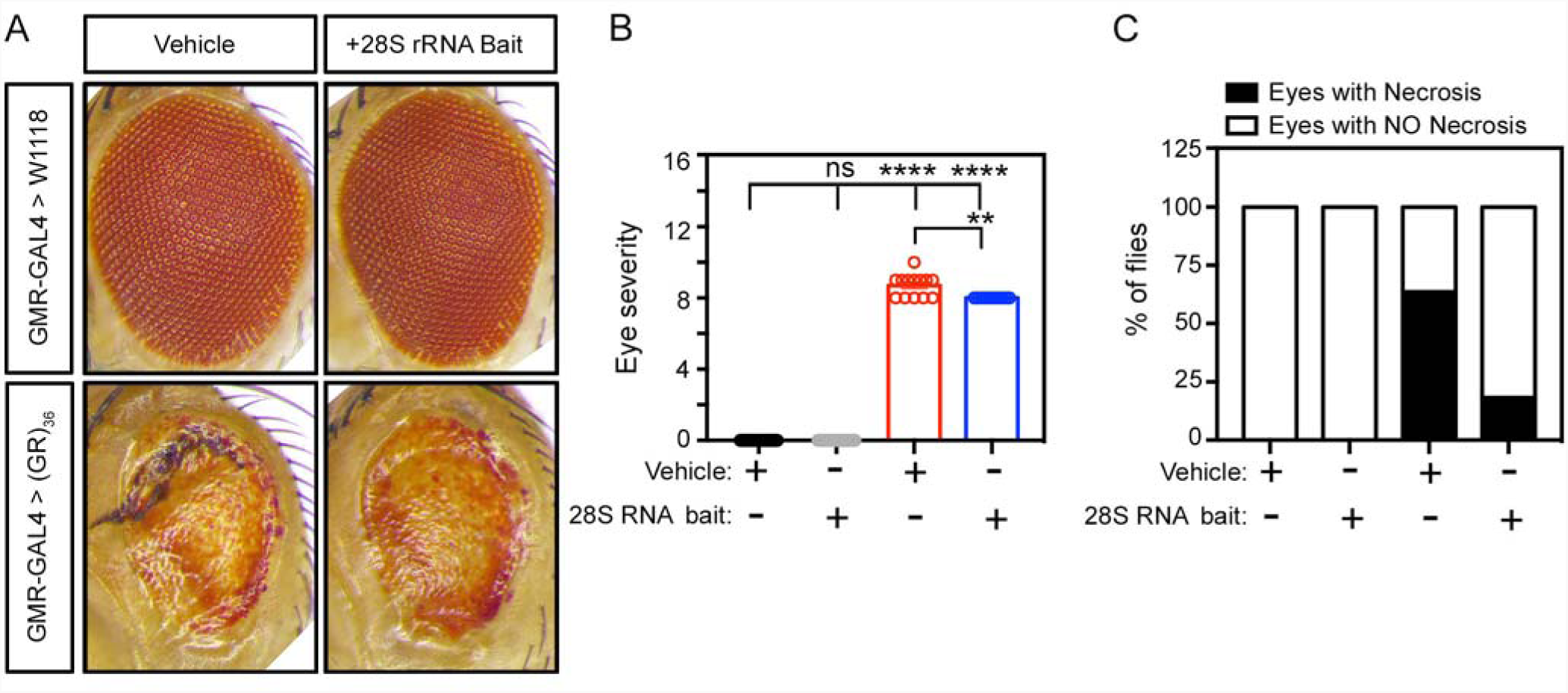
RNA-based strategy to inhibit the toxic effects of poly-GR *in vivo*. **A**. Representative eye images from (GMR-GAL4 x W1118) and (GMR-GAL4 x (GR)_36_) mutant flies treated or non-treated with 28S rRNA-based bait. **B**. Bar graph displaying the level of eye degeneration in the different conditions as referred to H. Each dot represents values of a single fly. ANOVA test, p**<0.01; p****<0.0001; ns, not significant. **C**. Graph bar showing the percentage of (GMR-GAL4 x W1118) and (GMR-GAL4 x (GR)_36_) flies with or without eye necrosis.

## DISCUSSION

The discovery of the C9-HRE as the most prevalent genetic driver of ALS/FTD has stimulated intense interest in deciphering the pathophysiology associated with this mutation. Several studies have shown that C9-DPR proteins have detrimental effects in cellular systems and model organisms (Choi et al., 2019; Freibaum et al., 2015; Hao et al., 2019; Hartmann et al., 2018; Jovicic et al., 2015; Kwon et al., 2014; Lee et al., 2016; Mizielinska et al., 2014; Tao et al., 2015; Wen et al., 2014; Zhang et al., 2018b; Zhang et al., 2019). We combined computational and experimental approaches to understand how the interaction of poly-GR with RNA contributes to toxicity. We found that poly-GR directly binds to multiple rRNA species in cells and impedes ribosomal homeostasis. We showcased the strength of the poly-GR/rRNA interaction by using a custom rRNA-based oligonucleotide, which reduced the malignant effects of poly-GR on rRNA levels and localization, protein translation and toxicity in multiple model systems. Our findings reinforce the importance of ribosomal impairment in C9-ALS/FTD and highlight a novel approach for protecting against poly-GR pathological mechanisms.

The characterization of the physicochemical features of poly-GR and poly-PR underscored a number of similarities, as well as critical structural differences that likely define their localization, molecular interaction profile and toxic potential. Although a recent computational study predicts that poly-GR and poly-PR form different oligomers containing what they named double-helix structures (Zheng et al., 2021), our analysis consistently shows that poly-GR acquires a random coiled conformation, while poly-PR is highly enriched in β-sheets. The discrepancies between Zheng et al., and our study are likely driven by the different computational methods. We used continuous MD (CMD), which is an unbiased method that models experimental conditions and accounts for the dynamics of the system. Zheng et al., used discontinuous MD, which favors interactions beyond any unbiased parameterization and thus tends to predict the formation of organized, ordered structures. Further, our computational data was validated by distinct empirical methods that confirmed the different secondary structures predicted for R-DPRs, where the higher rigidity of prolines compared to glycines play a critical role (Jafarinia et al., 2020). Interestingly, the secondary configuration of poly-PR confers a more stretched conformation, allowing more pronounced exposure of positive charges and a distinct adaptability to interact with complex molecular geometries such as the ones that are required during phase separation (Boeynaems et al., 2017; Flores et al., 2016; Jafarinia et al., 2020; Kanekura et al., 2018; Lee et al., 2016).

While the size of native DPR proteins produced in physiological models remains unknown, our analysis suggests that their secondary structural features are principally maintained irrespective of repeat number. This is due to the relatively short-range character of the secondary structures that are formed by the DPR amino acids. Several computational studies have demonstrated that sequences of 60 or less amino acids are sufficient for modelling the full conformational space of simple repetitive sequences (Cossio et al., 2010; Zamuner et al., 2015). It is noteworthy that the DPR proteins are unlikely to form a well-defined tertiary structure since at least four distinct amino acids are required for peptide sequences to attain a specific tertiary disposition (Cardelli et al., 2019; Nerattini et al., 2020). Thus, our findings support the notion that C9-HRE toxicity is threshold dependent and does not strongly correlate with repeat size (Cammack et al., 2019; Gendron et al., 2017; Gijselinck et al., 2016; Suh et al., 2015; van Blitterswijk et al., 2013). At the same time, our conclusions are tempered by the fact that DPRs can be heterogenous in content (McEachin et al., 2020; Tabet et al., 2018), while in patients the length of DPRs may be several hundred amino acids longer than the ones we investigated here, and size can affect their subcellular distribution and molecular interactions (Bennion Callister et al., 2016).

Our work strongly suggests that poly-GR compromises ribosomal homeostasis and impedes the ability of ribosomes to mediate protein translation. We specifically observed that poly-GR affected the subcellular distribution of rRNA, leading to a high N/C ratio within *in vitro* models, and patient tissue. We hypothesize that this effect is likely the result of poly-GR binding to multiple rRNA species found in both the nucleus and the cytoplasm. This finding is well-aligned with previously described defects in ribosomal biogenesis in the nucleus and protein translation in the cytosol (Hartmann et al., 2018; Kwon et al., 2014; Lee et al., 2016; Loveland et al., 2022; Moens et al., 2019; Wen et al., 2014; White et al., 2019; Zhang et al., 2018a). Moreover, a recent study demonstrated that C9-HRE patient-derived cells exhibit profound destabilization of ribosomal transcripts (Tank et al., 2018), and we propose that R-DPRs may be contributing to this defect. Importantly, the shift in rRNA that we observed could also be attributed to a previously described interaction of R-DPRs with nuclear pore proteins (Jovicic et al., 2015; Lee et al., 2016; Shi et al., 2017; Zhang et al., 2018b), although the effects of these interactions on N/C transport of rRNA-protein complexes remain unclear (Hayes et al., 2020; Vanneste et al., 2019; Zhang et al., 2018a).

Postmortem analysis of ALS/FTD patient tissue (Gomez-Deza et al., 2015; Mackenzie et al., 2015; Mori et al., 2013) and mouse models with C9-GR (Choi et al., 2019; Zhang et al., 2018b) has shown that the accumulation of poly-GR aggregates in the nucleus is a rare event. While one cannot discount the significance of soluble poly-GR that may exist in the nucleus and may be harder to detect by IHC, our findings provide additional insight to this observation. We found that the nuclear accumulation of poly-GR in human MNs is indeed a rare event (~10% of MNs overexpressing poly-GR exhibit nuclear aggregation), but that nuclear aggregation strongly predisposed MNs to degeneration. Thus, detecting surviving neurons with poly-GR nuclear aggregates postmortem would be unlikely.

CLIP-Seq analysis revealed that rRNA was the major RNA target of poly-GR in cells. While the potential for R-DPRs to interact with negatively charged molecules such as RNA had been established (Boeynaems et al., 2017; Boeynaems et al., 2019; Jafarinia et al., 2020; Kanekura et al., 2016; Lafarga et al., poly-GR can bind to both immature rRNA species that are found in the nucleolus, as well as fully processed rRNA found within cytosolic ribosomes. The preferential interaction with multiple rRNA species was unexpected, since the physicochemical properties of arginine-rich DPRs, as well as the biochemical assays *in vitro* suggest that they should exhibit rather promiscuous affinity for any available RNA molecule (Lafarga et al., 2021). However, one should consider that ribosomes are highly abundant in the cell and rRNA is topologically exposed all around the ribosomal surface along with RPs, many of which contain LCDs (Uversky, 2013) (Table S3). This would likely explain the recurrent precipitation of ribosomal proteins in multiple IP-MS studies (Hartmann et al., 2018; Lee et al., 2016; Lopez-Gonzalez et al., 2016; Radwan et al., 2020; Tao et al., 2015; Yin et al., 2017). Interestingly, our dynamic simulations predicted that it is the RNA that mediates the binding of R-DPRs to the ribosome, at least in the specific region modelled. This region likely represents a physiological binding site as it accommodates both exposed rRNA and ribosomal protein subunits shown to bind to poly-GR (Hartmann et al., 2018; Lee et al., 2016; Lopez-Gonzalez et al., 2016; Radwan et al., 2020; Tao et al., 2015; Yin et al., 2017). However, it is likely that poly-GR interacts with ribosomes at multiple locations. In fact, a recent study based on cryo-EM analysis describes the accumulation of short synthetic R-(DPRs)_20_ within the polypeptide tunnel of assembling ribosomes *in vitro* and suggests that this accumulation perturbs protein translation (Loveland et al., 2022). Altogether, diverse experimental approaches indicate that R-DPRs can bind to multiple ribosomal regions and at different maturation stages (Figure S4A).

Our findings showcase the importance of empirical data in cellular context in contrast to using *in vitro* experiments (i.e., purified DPRs with RNA in a test tube). We found that GR exhibited high affinity for tRNAs *in vitro*, while in live cells, although highly abundant, tRNAs are in fact de-enriched for GR binding. In further support of this argument, Balendra et al., recently performed similar CLIP-Seq experiments for poly-PR in cells and unexpectedly found that it binds specific mRNAs harboring a GAAGA sequence rather than exhibiting broad unspecific binding to all classes of RNAs (Balendra et al., 2022).

Research studies around the C9 mutation have highlighted the non-canonical translation of DPR proteins as a pathway that can be targeted therapeutically. While there are several efforts focused on identifying the molecular factors that specifically mediate the production of all DPR proteins (Cheng et al., 2018; Cheng et al., 2019; Green et al., 2017; Moens et al., 2019; Sonobe et al., 2018; Tabet et al., 2018; Westergard et al., 2019; Yamada et al., 2019), this may prove challenging. Alternatively, individual DPRs can be targeted by specific antibodies (Nguyen et al., 2020; Zhou et al., 2017), or as we propose here, by RNA oligonucleotides. The identification of a specific RNA target that natively interacts with poly-GR provided us with an opportunity to design a bait ribonucleotide molecule and assess its ability to protect cells by sequestering away poly-GR from its pathological interactions. Indeed, this molecule restored ribosomal homeostasis in (GR)_50_-transfected cells and ameliorated poly-GR toxicity in a *Drosophila* model and in iPSC-derived MNs overexpressing poly-GR or carrying *C9orf72* high-repeat expansions.

The mitigating effects of single strand RNA and DNA molecules have been recently explored in the context of other ALS/FTD model systems. Lafarga and colleagues showed that non-coding single strand DNA reduces the toxicity of synthetic R-DPRs treatments in U2OS cells and mouse motor neurons (Lafarga et al., transition of C9 R-DPRs (Boeynaems et al., 2017; Boeynaems et al., 2019), as well as to alleviate some of the pathophysiological mechanisms associated with R-DPR overexpression (Hayes et al., 2020; Lafarga et al., 2021). More aligned with the strategy that we propose here, recent studies have shown that short, chemically modified and sequence-specific RNA oligonucleotides of known TDP-43 targets can prevent inclusions and rescue mutant TDP-43 neurotoxicity (Mann et al., 2019). Although more work is required to understand how the binding of the 28S rRNA bait to poly-GR alleviates its pathophysiology, the promising results we present here support the notion that using bait RNAs is not only useful to study RNA-protein interactions (Jazurek et al., 2016), but also to protect neurons from the detrimental effects of mutant or aberrant proteins (Odeh and Shorter, 2020).

## ACKNOWLEDGMENTS

We are grateful to the following funding sources: US National Institutes of Health (NIH) National Institute of Neurological Disorders and Stroke (NINDS) and National Institute of Aging (NIA) R01NS104219 (E.K), NIH/NINDS grants R21NS107761 and R21NS107761-01A1 (E.K), the Les Turner ALS Foundation (E.K), the AFM-Telethon French Muscular Dystrophy Association Trampoline Grant #23648 (J.A.O), the Center for Regenerative Nanomedicine at the Simpson Querrey Institute (S.I.S and T.C), the New York Stem Cell Foundation (E.K), and the Intramural Research Program of the NIH, National Cancer Institute (NCI), Center for Cancer Research (M.B and S.L.W), the AFM-Telethon postdoctoral fellowship (J.A.O), the Ramon y Cajal fellowship RYC2019-026980-I (J.A.O), Gipuzkoa Foru Aldundia 2019-FELL-000017-01 and Maria de Maeztu Units of Excellence MDM-2017-0720 (I.R.S), NINDS grants R01NS097850 and R01NS131409 (J.K.I.), the New York Stem Cell Foundation (J.K.I.), Department of Defense grants PR211919 and W81XWH2110131 (J.K.I.), and the John Douglas French Alzheimer’s Foundation (J.K.I.). J.K.I. is a New York Stem Cell Foundation – Robertson Investigator and the John Douglas French Alzheimer’s Foundation Endowed Associate Professor of Stem Cell Biology and Regenerative Medicine. E.K is a Les Turner ALS Center Investigator and a New York Stem Cell Foundation – Robertson Investigator.

## CONFLICTS STATEMENT

J.K.I. is a co-founder of AcuraStem Inc, and Modulo Bio, a scientific advisory board member of Spinogenix, Vesalius Therapeutics, and Synapticure, and a paid employee of BioMarin Pharmaceutical Inc. None of these companies were involved in this work.

## METHODS

### All-atoms molecular dynamics simulations

Peptide structures were built in Avogadro (Hanwell et al., 2012) and the ribosome structure was obtained from *Oryctolagus cuniculus* (PDB ID: 5LZS) (Shao et al., 2016). All simulations were carried out in GROMACS 5.0.4. (Abraham et al., 2015) using Chemistry at Harvard Macromolecular Mechanics (CHARMM) 36 force field for all-atomistic dynamic simulations that represent all the atoms of the system (Best et al., 2012; Brooks et al., 2009; Denning et al., 2011; MacKerell et al., 1998; MacKerell et al., 2000; Vanommeslaeghe et al., 2010). Force field has been widely used for simulation studies of proteins, peptides and nucleic acids (Babaian et al., 2020; Cook et al., 2020; Kognole and MacKerell, 2020; Suomivuori et al., 2020; Zhu et al., 2012). The experimental system was set up with a constant number of molecules, pressure and temperature, in a 100 mM NaCl environment with TIP3P water (Jorgensen et al., 1983). The simulation box size was set up allowing a margin of 2 nm at each side of the ribosome or of stretched DPR. DPRs simulations were done for 60 ns. Ribosome was simulated for 100 ns. Combined DPR-ribosome systems were set up placing the equilibrated DPR of 50 repeats close to the area of interest of the equilibrated ribosome and simulated for 60 ns. Molecular projections do not show the backbone nor the hydrogens which allows a better visualization of the molecular conformation and side chains. However, the multiple calculations considered all the molecular components. All visualizations were rendered using Visual Molecular Dynamics (VMD) package (Humphrey et al., 1996). The following analyses were also carried out using GROMCAS 5.0.4: R_g_ (radius of gyration) ≈ peptide extension ≈ 1/folding SASA (solvent accessible surface area): Measure the exposure of the molecule to the solvent. AP = SASA(0)/SASA(i); is a normalization to show the tendency of the distinct DPRs to aggregate (AP=1. means fully soluble, AP>1 means aggregated).

RMSD (root mean square deviation) measures deviations in the structure with time relative to a reference structure (the initial conformation). Higher RMSD values involve bigger differences with respect to the initial structure. Constant RMSD means stable structures and, hence, reaching a plateau is indicative of equilibrated simulations. RMSF (root mean square fluctuations) measures the fluctuations of atoms in nanometers respect to a reference structure in a given time range. Measurements through 10 ns in the equilibrated region gives a value of the mobility of atoms. Lower RMSF involves lower mobility and, hence, higher stability of the selected atoms in the reference position. <u>Simulation Procedure:</u> All the systems were minimized using steepest decent for 50000 steps or until forces on atoms converged below 1000 pN. The systems were equilibrated in NVT ensemble at 300 K for 100 ps, and then in NPT at 300 K and 1 atm for 1 ns, adding constraints in backbone atoms. Simulations were then run for the specified time (100 ns ribosomes and 60 ns DPRs and DPR-ribosome) systems in NPT ensemble. Equilibrations and simulations used 2 fs timestep and periodic boundary conditions in the three spatial coordinates. Verlet cut-off scheme was employed for non-bonded interactions with a cut-off radius of 1.2 nm (shifting van der Waals to zero from 1.0 nm) (Verlet, 1967) and Particle Mesh Ewald for long-range electrostatics (Darden et al., 1993). Temperature was controlled using velocity rescaling algorithm (τ_T_ =0.1 ps) (Bussi et al., 2007). Pressure was kept constant using Berendsen algorithm for the NPT equilibration (Berendsen et al., 1984) and Parrinello-Rahman in the simulations (both τ_P_=2 ps)(Parrinello and Rahman, 1981).

### Coarse-grained molecular dynamics simulations

Aggregation propensities (AP) of the different DPRs were calculated from coarse-grained molecular dynamics simulations using the MARTINI force field (version 2.2) with coil input secondary structure (de Jong et al., 2013; Marrink et al., 2007; Monticelli et al., 2008). This model maps up to 4 heavy atoms to 1 bead in order to speed up the simulations. This force field has been previously employed to measure peptide AP (Frederix et al., 2015; Frederix et al., 2011). The results for each DPR is the average of two independent simulations at different simulation box volumes to minimize any finite size effect in the results. Systems were built with constant number of amino acids 1200 or 2400 in a cubic box of 17.1 nm or 21.6 nm of side, respectively. Final concentrations are 22, 11, 7 and 4 mM for 10, 20, 30 and 50 number of repeats, respectively. Simulation procedure: All systems were minimized using steepest descent for 5000 steps or until forces converged below 200 pN. The systems were equilibrated in NPT ensemble at 303 K and 1 atm for 1000 steps using sequentially 1, 5, 10 and 20 fs timestep. Aggregation simulations were then run for 5 µs in the same ensemble with periodic boundary conditions in the three spatial coordinates. A 1.1 nm cut-off was applied for non-bonded interactions using potential-shift for Lennard-Jones and reaction field, with a dielectric constant of 15, for electrostatics (de Jong et al., 2016). V-rescale and Berendsen algorithms were used to keep temperature (τ_T_=1.0 ps) and pressure (τ_P_ =6 ps), respectively, constant (Bussi et al., 2007).

### Peptide synthesis

Peptides were synthesized via standard 9-fluorenyl methoxycarbonyl (Fmoc) solid-phase peptide chemistry on Wang resin using a CEM Liberty Blue automated microwave peptide synthesizer. Automated coupling reactions were performed using 4 eq. of Fmoc-protected amino acid, 4 eq. of N,N‘-diisopropylcarbodiimide (DIC), and 8 eq. of ethyl(hydroxyimino)cyanoacetate (Oxyma pure) and removal of Fmoc groups was achieved with 20% 4-methylpiperidine in DMF. Peptides were cleaved from the resin using standard solutions of 95% TFA, 2.5% water, 2.5% triisopropylsilane (TIS) and then precipitated with cold ether to yield the crude peptide product. The crude product was purified by preparative reverse-phase high-performance liquid chromatography (RP-HPLC) using a Phenomenex Kinetex column (C_18_ stationary phase, 5 μm 100 Å pore size, 30 × 150 mm) on a Shimadzu model prominence modular HPLC system equipped with a DGU-20A5R degassing unit, two LC-20AP solvent delivery units, a SPD-M20A diode array detector and a FRC-10A fraction collector, using H_2_O/CH_3_CN gradient containing 0.1% CF_3_COOH (v/v) as an eluent at a flow rate of 25.0 mL/min.

### Liquid Chromatography-Mass Spectrometry (LC-MS)

Analytical RP-HPLC was performed at 40 °C using a Phenomenex Jupiter 4 µm Proteo 90 Å column (C12 stationary phase, 4 µm, 90 Å pore size, 1 × 150 mm) on an Agilent model 1200 Infinity Series binary LC gradient system, using H_2_O/CH_3_CN gradient containing 0.1% CF_3_COOH (v/v) as an eluent at a flow rate of 50 µL/min. Electrospray ionization mass (ESI-mass) spectrometry was performed in positive scan mode on an Agilent model 6510 Quadrupole Time-of-Flight LC/MS spectrometer using direct injection.

### Optical density measurements

Optical density (O.D.) measurements were performed on a BioTek model Cytation 3 cell imaging multi-mode reader.

### Circular dichroism (CD)

CD spectra were recorded in a Jasco J-815 spectropolarimeter using quartz cells of 100 µm pathlength. Spectra were background subtracted and are the average of three scans using continuous scanning mode at a speed of 100 nm/min and standard sensitivity. Final spectra are normalized to concentration. Further spectra analysis was carried out employing the server BESTSEL (https://bestsel.elte.hu/) (Micsonai et al., 2018; Micsonai et al., 2015). Other CD analysis tools were considered but they were discarded because, unlike BESTSEL, they require the protein family and/or type as input. It is noteworthy that these methods are strongly protein-type and parameter-dependent, benchmarked with crystal information and thus form relatively highly order proteins. Additionally, none of the benchmarking sets are expected to have such high percentage of glycine and/or proline as R-DPRs, which can substantially affect the signal. Therefore, the deconvolution results obtained must be taken with caution and only the correlation with other techniques can validate their interpretation. Additionally, the spectra of three reference systems are included. These are a strong β sheet forming protein, as well as an α helix and disorder peptide, which are, respectively, UnaG (PDB:4I3B, 10 µM), consensus tetratricopeptide repeat protein (PDB: 7OBI, 5 µM), and the peptide AAGGEE (50 µM), capped in the N-terminus with a palmitic acid and in the C-terminus by an amide.

### Fourier-transform infrared spectroscopy (FTIR)

FTIR spectra were recorded on a Bruker Tensor 37 FTIR spectrometer. Spectra shown are the average of 25 scans with a resolution of 1 cm^−1^. Samples were prepared in deuterated water (D_2_O) to displace its vibrations from the region of interest. Liquid samples were placed between two CaF_2_ windows with 50 µm pathlength and background subtracted using the solvent. Solid FTIR was measured using attenuated total reflectance (ATR) module on lyophilized samples and using background subtraction to remove signals from atmospheric H_2_O and CO_2_. The liquid FT-IR spectra were deconvoluted using Gaussian lineshape to better illustrate the contributions of the different modes (Yang et al., 2015).

### RNA-dipeptide repeats binding assay

The RNA solutions were diluted with HEPES buffer ([phosphate] = 200 µM) and incubated for 30 min at 25 °C. A HEPES buffer solution of dipeptide repeats (10 mM) was then added to the RNA solutions, and the mixture was pipetted 30 times. 50 µL of these suspensions were put into triplicate wells of a 96-well plate, and their optical density at 600 nm were recorded.

### Estimation of the content of phosphates in the RNA sample

The content of phosphates in the RNA sample was estimated based on assumptions that the RNA is 100% pure and its counter cation is sodium. Molecular weight of the RNA repeating unit (343.43 g/mol) was calculated by averaging the molecular weights of adenine (351.19 g/mol), guanine (367.19 g/mol), cytosine (327.17 g/mol) and uracil (328.15 g/mol). As the concentration of RNA was shown to be 5.4 µg/µL, the concentration of phosphates was therefore estimated to be 15.7 mM.

### Cell culture models

#### HEK-293FT models

HEK-293FT cells were grown in DMEM (Corning) supplemented with Glutamax (Gibco) and 10% fetal bovine serum (FBS, Gibco). HEK-293FT cells were dissociated by incubating for 5 min with Trypsin-EDTA (Gibco) at 37°C. All cell cultures were maintained at 37C and 5% CO_2_ and tested for mycoplasma monthly.

#### iPSC models

The induced pluripotent stem cell line (18a: female, 48 years old) derived by retroviral transduction of skin fibroblasts from a healthy control individual (Boulting et al., 2011), was utilized to generate human MNs as described previously (Ziller et al., 2018). iPSCs were maintained on Matrigel (BD Biosciences) with mTeSR1 media (Stem Cell Technologies) and passaged on a weekly basis using 1mM EDTA or Accutase (Sigma). At 70% confluency, iPSC cultures were dissociated using Accutase and plated at a density of 10^5^cells/cm^2^ with 10µM ROCK inhibitor (Y-27632, DNSK International) in mTeSR1. Next day (day 0), media was replaced with N2B27 medium (50% DMEM:F12, 50% Neurobasal, supplemented with non-essential amino acids (NEAA), Glutamax, N2 and B27; Gibco, Life Technologies) containing 10µM SB431542 (DNSK International), 100nM LDN-193189 (DNSK International), 1 µM Retinoic Acid (RA, Sigma) and 1µM of Smoothened-Agonist (SAG, DNSK International). The culture medium was changed daily until day 6, then switched to N2B27 medium supplemented with 1µM RA, 1µM SAG, 5µM DAPT (DNSK International) and 4µM SU5402 (DNSK International). Cells were fed daily until day 14, when MNs were dissociated using TrypLE Express (Gibco, Life Technologies) supplemented with DNase I (Worthington) and plated at a density of 10,000 cells /well in Incucyte® Imagelock 96-well Plate (Sartorius) pre-cultured with 20,000 mouse glial cells/well. Co-cultures were feed 3 times a week with NBM medium (Neurobasal, NEAA, Glutamax, N2 and B27) supplemented with 1% FBS, Ascorbic acid (0.2 µg/ml; Sigma-Aldrich), BDNF, CNTF and GDNF (10ng/mL, R&D systems).

#### iMNs models

Human lymphocytes from healthy subjects and ALS patients were obtained from the NINDS Biorepository at the Coriell Institute for Medical Research and reprogrammed into iPSCs using episomal plasmids as previously described (Okita et al., 2011; Shi et al., 2019; Shi et al., 2018). iPSCs were differentiated into fibroblast-like cells to enable efficient retroviral transduction as described previously (Shi et al., 2019). Reprogramming of fibroblast-like cells was performed in 96-well plates (5 x 10^3^ cells/ well) or 13-mm plastic coverslip (3 x 10^4^ cells/ coverslip) that had been pre-coated with 0.1% gelatin (1 hour, room temperature) and laminin (4°C, overnight). Retroviruses encoding seven iMN factors (*Ngn2*, *Lhx3*, *Isl1*, *NeuroD1*, *Ascl1*, *Brn2* and *Myt1l*) were added in 150 μL of fibroblast medium (DMEM plus 10% FBS) per 96-well with 8 μg/ml polybrene. Hb9::RFP lentivirus were added to the cultures 24 hours post-transduction with seven iMN factors. On day 4, primary mouse cortical glial isolated from postnatal day 3 ICR pups were added to the transduced cultures in MEM (Life Technologies) and 10% donor equine serum (HyClone). On day 5, cultures were switched to N3 medium containing DMEM/F12 (Life Technologies), 2% FBS, 1% penicillin/streptomycin, glutamax, N2 and B27 supplements (Life Technologies), 7.5 μM RepSox (Selleck), and 10 ng/ml each of FGF, GDNF, BDNF, and CNTF (R&D). The cultures were maintained in N3 with neurotrophic factors (RepSox, FGF, GDNF, BDNF, and CNTF) and changed every other day.

### Donor information and karyotyping results for iPSCs

| Line name | Disease status | Gender | Age of onset | Age at sampling | Race | iPSC karyotype |
| --- | --- | --- | --- | --- | --- | --- |
| 18a | Control | Female | Not available | 48 | Caucasian | Normal |
| ND03231 | Control | Male | Not available | 56 | Caucasian | Normal |
| ND03719 | Control | Male | Not available | 33 | Caucasian | Normal |
| ND05280 | Control | Female | Not available | 72 | Caucasian | Normal |
| ND06769 | *C9ORF72*<br>ALS | Female | 45 | 46 | Caucasian | Normal |
| ND10689 | *C9ORF72*<br>ALS | Female | 49 | 51 | Caucasian | Normal |
| ND12099 | *C9ORF72*<br>ALS | Male | 48 | 49 | Caucasian | Normal |

### DPR overexpression in cell culture models

For overexpression experiments, 40% confluent HEK-293 cells were transfected with HilyMax transfection reagent (Dojindo Molecular Technologies) according to manufacturer guidelines. Briefly, DNA was mixed with HilyMax (1µg DNA:3µL HilyMax ratio) in Opti-MEM medium (Gibco) and incubated for 15 min at room temperature (RT) before being added to cells. Cells were incubated with transfection mixture for 4hrs at 37°C and then media was replaced. Analyses made on transfected cells were performed 48 to 72 hrs after transfection.

Overexpression of GFP or GFP-(GR)x50 in iPSC-derived MNs was achieved through lentiviral infection. MNs were infected with lentiviral stocks for 24 hours on day 17 of differentiation. We performed real-time quantitative live-cell imaging analysis using the Incucyte® Live-Cell Analysis System as well as immunocytochemistry at multiple time points.

### iMN survival assay

Transfection was performed on day 15 of the iMN conversion assay. Next day, a complete medium change without RepSox, FGF, GDNF, BDNF, or CNTF was done and longitudinal tracking of iMNs was initiated. Images of *Hb9*::RFP+ iMNs were taken using Molecular Devices ImageExpress once every other day for 14 days. Tracking of neuronal survival was performed using SVcell 3.0 (DRVision Technologies) or ImageJ. Neurons were scored as dead when their soma was no longer detectable by RFP fluorescence.

### Plasmids and virus stock preparation

HEK-293 cells were transfected with the pcDNA3.1 plasmid containing GFP, GFP-(GP)x10-, or GFP-(GR)x50, in which alternative codons were used to generate DPRs without generating the (GGGGCC)n transcript. These constructs were made and kindly shared by Petrucelli’s lab (Zhang et al., 2014). iPSC-derived MNs were transduced with lentivirus containing the above-mentioned GFP or GFP-(GR)x50 sequences inserted in the lentivector backbone CD510B. Packaging from lentiviral constructs was performed in 293FT cells by transfecting packaging components contained in pMD2.G and psPAX2 vectors with HilyMax (Dojindo Molecular Technologies). Virus was collected from cell media 48 to 96 hours after transfection. Lentiviral concentrated stocks were obtained after filtering collecting media and centrifugation at 25,000 g for 2h at 4°C. For iMNs models, the complementary DNA (cDNA) for each iMN factor (*Ngn2, Lhx3, Isl1, NeuroD1, Ascl1, Brn2* and *Myt1l*) was purchased from Addgene and cloned into the pMXs retroviral expression vector using Gateway cloning technology (Invitrogen). The Hb9::RFP lentiviral vector and rtTA3 lentiviral vector were also purchased from Addgene (ID: 37081; 61472). Viruses were produced as follows. HEK293T cells were transfected in a 10-cm dish at 80-90% confluence with viral vectors containing each iMN factor and viral packaging plasmids (PIK-MLV-gp and pHDM for retrovirus, pPAX2 and VSVG for lentivirus) using polyethylenimine (PEI)(Sigma-Aldrich). The medium was changed 24 hours after transfection. Viruses were harvested at 48 hours and 72 hours after transfection. Viral supernatants were filtered with 0.45-μm filters, incubated with Lenti-X concentrator (Clontech) for 24 hours at 4°C, and centrifuged at 1,500 g at 4°C for 45 minutes. Pellets were resuspended in DMEM plus 10% FBS (200 μL per 10-cm dish of HEK293T) and stored at −80°C.

### CLIP-Seq

CLIP experiments were performed as described (Huppertz et al., 2014). Briefly, cells were crosslinked in a Spectroline UV crosslinker using 100 mJ/cm^2^ UVC (254 nm). Cells were lysed in 50 mM Tris-HCl pH 7.4, 150 mM NaCl, 1% NP-40, 0.5% sodium deoxycholate, 0.1% SDS with EDTA-free complete mini protease inhibitor cocktail (Roche) and 1 mM phenylmethylsulfonyl fluoride (PMSF). After sonicating on ice using a Branson sonifier, lysates were incubated with 4 U/ml Turbo DNase (Invitrogen) and 0.2 to 0.002 units/ml of RNase I (Invitrogen). Immunoprecipitation was performed by incubating the lysates with GFP-Trap Magnetic beads (Chromotek cat: gtd-10) for 1.5 hours at 4°C. Afterwards, beads were washed twice with high salt buffer (50 mM Tris-HCl pH 7.4, 1 M NaCl, 1% NP-40, 0.5% sodium deoxycholate, 0.1% SDS) and twice with PNK buffer (20 mM Tris-HCl pH 7.4, 10 mM MgCl_2_, 0.2% Tween-20). RNA 3’ ends were dephosphorylated with PNK as described (Huppertz et al., 2014), followed by two washes each with high salt and PNK buffers. 3’ linker ligation was performed on beads overnight at 16°C followed by two additional washes each with high salt and PNK buffer. After labeling with [γ-^32^P]-ATP and T4 RNA ligase, RNPs were resolved on NuPAGE 4-12% Bis-Tris gels (Invitrogen) run in NuPAGE MOPS-SDS running buffer (Invitrogen). After gel fractionation, RNPs were transferred to Amersham Protran 0.45 uM nitrocellulose membranes (Cytiva) and labeled complexes visualized using a storage phosphor screen and developed on a Typhoon FLA 9000 scanner. RNA-protein complexes of the expected size were excised from the nitrocellulose membranes, along with the same size region from GFP, GFP-(GP)_10_, or untransfected control experiments. Associated proteins were removed by digesting with 1 mg/ml proteinase K (Invitrogen) for 20 minutes at 37°C in PK buffer (100 mM Tris-HCl pH7.4, 50 mM NaCl, 10 mM EDTA) followed by a second digestion for 20 minutes at 37°C in the presence of 3.5 M urea. Afterwards, RNA was extracted with phenol/chloroform and reverse transcription performed using Superscript III (Invitrogen) using the following primers: library 1 GPF - Rt1clip, GFP-(GR)_50_ - Rt6clip; library 2 untransfected - Rt1clip, GFP-(GP)10 - Rt6clip, GFP-(GR)_50_ - Rt9clip. cDNA was size selected and circularized with Circligase II (Epicentre). Circularized cDNA was cut with BamHI and amplified using Accuprime Supermix I (Invitrogen). The PCR cycle number was optimized to prevent overamplification of the library. Amplified samples were sequenced on the Illumina MiSeq platform in single end read mode with 110 nt reads.

### CLIP-Seq bioinformatics

For mapping to the whole genome, barcoded sequencing libraries were demultiplexed allowing 1 nt barcode mismatch, and adapter sequences and low-quality bases were filtered using iCount (version 2.0) (https://github.com/tomazc/iCount). Trimmed, single-end reads were aligned to the hg38 genome (Gencode V32) using Novoalign (version 4.03.03). Parameters for alignment included: score threshold (-t) set to 15,3, the minimum number of bases for alignment (-l) set to 20, the gap penalty (-x) set to 4, gap opening penalty (-g) set to 20, trimming step size (-s) set to 1, score difference (-R) set to 0 and multimapping reporting (-R) set to ALL. The quality of the sequenced libraries was assessed per sample using FastQC (version 0.11.5) https://www.bioinformatics.babraham.ac.uk/projects/fastqc/, FastQ Screen (version 2) (Wingett and Andrews, 2018) and samtools (version 1.13) (Li et al., 2009). Samples were deduplicated using UMI-tools (Smith et al., 2017), with multi-mapping detection. On average 5.33 unique UMIs were detected per position.

To determine the location of CLIP peaks, overlapping reads were collapsed to create a list of genomic regions using bedtools (version 2.30.0) (Quinlan and Hall, 2010). Peaks found within 50 nt of one another were combined to a single feature. Reads associated with CLIP peaks were counted using FeatureCounts (version 2.0.2) (Liao et al., 2014) with the following parameters: the number of reads supporting each exon-exon junction was included, the minimum number of overlapping bases in a read required for assignment as 1, and strand specific read counting was performed. Analysis was performed without multi-mapped reads (unique reads only) and with multi-mapped reads included (unique + multi-mapped reads). Multi-mapped reads were assigned a fractional count of 1/x, where x is the total number of alignments reported for the same read. CLIP peaks with > 5 unique + fractional multi-mapped reads were annotated with overlapping genomic features. Protein coding and ncRNA features were identified from the Gencode hg38 V32 while ncRNA and repeat regions were identified from hg38/GRCh38 Repeatmasker annotations and rRNA annotations were identified from RefSeq GRCh38.p13 (GCF_000001405.39). All features annotated for each peak are reported in Table S2. Representative peaks with multiple annotations were initially examined manually to determine the correctness of the annotations. Based on this analysis, preference was given to annotations as follows: rRNA > ncRNA > protein coding: exon > repeat element > pseudogene > lncRNA: exon > antisense feature > protein coding: intronic > lncRNA: intronic > no feature.

To identify specific sites of RNA binding, MAnorm (version 1.1.4) (Shao et al., 2012) was used to identify enriched CLIP peaks in GFP-(GR)_50_ vs GFP samples. Default parameters were used except for the following: shift size for both inputs (--s1,--s2,) was set to 0 to keep the peak binding site at the 5’ end and the summit-to-summit distance cutoff for common peaks (-d) was set to 25 to ensure only overlapping peaks between samples were compared. 10,000 simulations (-n) were performed to test enrichment. CLIP peaks with a fold-change > 2 and a p-value <.05 were identified as significant binding events.

For mapping to the RNA45SN1 locus, PCR duplicates were removed by collapsing identical sequences using the FastX_collapser. PhiX spike in control reads were removed by mapping to Coliphage phi-X174, complete genome (NCBI Reference Sequence: NC_001422.1). The remaining reads were demultiplexed based on barcodes using FastX_splitter, then adapter and UMI sequences were removed with fastX_trimmer and fastX_clipper, respectively (Fastxtoolkit version 0.0.14) (http://hannonlab.cshl.edu/fastx_toolkit/). Bowtie2-build was used to generate a genomic index from the RNA45SN1 (NC_000021.9) sequence. Reads were mapped to this index using Bowtie2 (version 2.4.4) (http://bowtie-bio.sourceforge.net/bowtie2/index.shtml) with the --very-sensitive-local option. Unmapped reads were extracted using Samtools-view and mapped to the hg38 human genome using STAR. Reads mapping to 45S were counted based on the regions with which they overlapped. Bedtools intersect was used to count reads in the various regions. For precursor regions (5’ETS, ITS1, ITS2 and 3’ETS) reads with at least one nucleotide in these regions were counted. For mature rRNA regions (18S, 5.8S and 28S) only reads mapping entirely within these regions were counted. The percentage of reads mapping to each region were represented as a percentage of the total collapsed read number.

To compare our rRNA reads to those present in other iCLIP datasets, fastq files for YTHDC2 iCLIP (SRR3175596) and HNRNPA1 iCLIP (ERR908337) were obtained from the NCBI Sequence Read Archive (SRA). Fastq files were mapped to the hg38 genome using STAR and demultimplexed using UMItools. Bedgraph files of the reads mapping to rRNA at chr21:8,436,754-8,446,360 were generated using Bedtools and normalized by the total number of reads mapping to this region in each library.

### RNA immunoprecipitations (RIP) followed by detection of RNAs by northern blot (NB) and RT-qPCR

HEK-293 cells expressing GFP- or GFP-(GR)_50_ were lysed in NET-2 (50mM Tris pH 7.5, 150 mM NaCl, 2.5 mM MgCl2, 0.5% NP-40) and 1 mM phenylmethylsulfonyl fluoride (PMSF). After sonicating in a Bioruptor Plus on high (30 seconds on, 30 seconds off) for 1 minute at 4 C, lysates were sedimented at 16,000 x g for 10 minutes. Cleared lysates were incubated with 25 μl of GFP-Trap magnetic beads for 1.5 hours at 4°C with rotation. Beads were washed twice in NET-2, transferred to a fresh tube, and washed two additional times. Beads were resuspended in 400 ul NET-2 and extracted with an equal volume of acid-phenol:chloroform (Invitrogen AM9722). After centrifugation at 16,000 x g for 15 minutes, RNA was precipitated from the aqueous phase by adding 1/10^th^ volume sodium acetate and 2.5 volumes 100% ethanol. After precipitation, RNA was fractionated in a 5% polyacrylamide/7M urea gel (to detect RNAs of less than 500 nts) or an 0.8% agarose/formaldehyde gel using the Tricine/Triethanolamine buffer system described by (Mansour and Pestov, 2013) to detect larger RNAs. RNA was transferred from polyacrylamide gels to Hybond-N (Cytiva) in 0.5X TBE for 16 hours at 150 mA. RNA was transferred from agarose gels to Hybond-N by capillary transfer overnight using 10X SSC (1.5 M NaCl, 150 mM sodium citrate pH 7). RNA was crosslinked to membranes using a Spectroline UV crosslinker and hybridized in modified (Church and Gilbert, 1984b) hybridization buffer (1% BSA, 2 mM EDTA, 200 mM NaHPO4 pH 7.2, 15% DI formamide, 7% SDS) using 5’-^32^P labeled oligonucleotides at 28°C. Northern probe sequences used in this study:

5.8S – GTGTCGATGATCAATGTGTCCTGCAATTCA

18S – CGCTCCACCAACTAAGAACG

28S – CCTGGTTAGTTTCTTCTCCTCC

7SL – CCATATTGATGCCGAACTTAGTGC

RPPH1 – CTGTTCCAAGCTCCGGCAAA and AATGGGCGGAGGAGAGTAGT

For RT-qPCR analyses, the RNA was reverse transcribed using the iSCRIPT cDNA Synthesis Kit (Bio-Rad) and qPCR was performed using iTaq Universal SYBR Green Supermix (biorad). Samples were run on a Bio-Rad CFX96 Real Time PCR System and analyzed using Maestro software (Bio-Rad).

Primer sequences used in this study:

28S_F – GGAGGAGAAGAAACTAACCAGG

28S_R – GTCTTCCGTACGCCACATGTC

18S_F – CTCAACACGGGAAACCTCAC

18S_R – CGCTCCACCAACTAAGAACG

5’ ETS_F – TCTAGCGATCTGAGAGGCGT

5’ ETS_R – CAGCGCTACCATAACGGAGG

ITS1_F – CAACCCCCTCTCCTCTTGGG

ITS1_R – GAGGTCGATTTGGCGAGGG

Y4_F – GGCTGGTCCGATGGTAGTGG

Y4_R – AAAGCCAGTCAAATTTAGCAGTGGG

Y5_F – AGTTGGTCCGAGTGTTGTGGG

Y5_R – AAAACAGCAAGCTAGTCAAGCGCG

tRNA-Glu_F – TCCCACATGGTCTAGCGG

tRNA-Glu_R – TTCCCACACCGGGAGT

### Immunoprecipitation (IP) followed by northern blot (NB) to detect bait RNA

GFP-GR_50_- and bait-transfected HEK293 cells were lysed with NET-2 buffer (50 mM Tris, 150 mM NaCl, 2.5 mM MgCl2, 0.5% NP-40) supplemented with protease inhibitors. Samples were sonicated at 4°C using Bioruptor Pico (Diagenode B01080010) on high mode, 30 seconds on and 30 seconds off for 2 cycles. Lysates were cleared by centrifugation at 16,000g for 10 minutes at 4°C. Supernatants were incubated with GFP-Trap magnetic beads (Chromotek gtd-10) at 4°C for 90 minutes using manufacturer recommended guidelines. The bead-sample complex was washed twice with NET-2, moved to a new tube, and washed two more times. Beads were resuspended in 400 μl NET-2 and extracted with an equal volume of phenol:chloroform:isoamyl 25:24:1 (Invitrogen 15593031). After centrifugation at 16,000 x g for 15 minutes, RNA was precipitated from the aqueous phase by adding sodium acetate to 0.5 M and 2.5 volumes 100% ethanol and incubating overnight at −20°C. After precipitation, RNA was fractionated in an 8% polyacrylamide/7M urea gel. RNA was transferred from the polyacrylamide gel to Hybond-N (Cytiva) in 0.5X TBE for 1.5 hours at 250 mA at 4°C. RNA crosslinking to Hybond-N was performed as described (Pall and Hamilton, 2008), incubating the membrane with chemical crosslinker at 60°C for 1 hour. Hybridization was performed in 1% BSA, 1 mM EDTA, 500 mM NaHPO4 pH 7.2, 7% SDS (Church and Gilbert, 1984a) using ^32^P-labeled oligonucleotides at 28°C.

### Immunoprecipitation (IP) followed by western blot (WB) or fluorescence intensity analysis

Cells were harvested in IP buffer (10 mM Hepes pH 7.6, 100 mM NaCl, 1 mM EDTA, 1 mM NaF, 2 mM Na_3_VO_4_, 1 mM DTT, 1 mM PMSF, 1% sodium deoxycholate, 10% glycerol, 0.1% SDS, 1% Triton X-100 and 1x protease inhibitor cocktail). Lysates were sonicated and protein concentrations determined with a BCA kit (Pierce). GFP and GFP-tagged proteins were immunoprecipitated from 1 mg of protein/sample with anti-GFP antibody (Abcam). Immunoprecipitation of the target antigen was performed using Dynabeads® Protein A (Novex, life technologies) following the manufacturer’s protocol (https://www.thermofisher.com/document-connect/document-connect.html?url=https%3A%2F%2Fassets.thermofisher.com%2FTFS-Assets%2FLSG%2Fmanuals%2FDynabeadsProteinA_man.pdf&title=RHluYWJlYWRzIFByb3RlaW4gQQ==). Eluted proteins were separated by SDS-PAGE followed by electrotransfer to a nitrocellulose membrane (Bio-Rad). The membranes were blocked in Tris-buffered saline (TBS, 50mM Tris, 150mM NaCl, HCl to pH 7.6) + 0.1% Tween 20 (Bio-Rad) + 5% non-fat dry milk (LabScientific) and incubated overnight at 4°C with primary antibodies: GAPDH (rabbit, 1:1000, Cell Signaling), GFP (goat, 1:1000, Abcam), RPL7A (rabbit, 1:1000, Cell Signaling Technology). Primary antibodies were diluted in TBS + 0.1% Tween + 5% BSA (Calbiochem). After several washes in TBS + 0.1% Tween, membranes were incubated with their corresponding secondary HRP-conjugated antibodies (1:5000, LI-COR Biotechnology). Protein signals were detected by a ChemiDocTM XRS+ (Bio-Rad), using the SuperSignal West Pico chemiluminescent system (Thermo Scientific).

For testing the interaction of Cy3-labeled rRNA baits with GFP *vs*. GFP-(GR)_50_, we performed anti-GFP IP using trap-bead GFP nanobodies (Chromotek, Proteintech) in GFP- and GFP-(GR)_50_-transfected cell substrates. Cy3 dye signal in the respective input cell substrates and eluted products that result from the magnetic-based IP was analyzed in a Cytation^TM^ 3 automated fluorescence plate reader (BioTek).

### Immunocytochemistry

Cells were fixed with 4% paraformaldehyde for 20 min, washed with PBS and permeabilized/blocked for 1h in PBS containing 10% normal donkey serum (Jackson ImmunoResearch) and 0.2% Triton. Samples were then incubated overnight at 4°C with primary antibodies: puromycin (mouse, 1:5000, Millipore), rRNA (mouse, 1:1000, Novus Biologicals), fibrillarin (rabbit, 1:2000, Abcam), NPM1 (mouse, 1:500, Santa Cruz Biotechnology). The next day, PBS + 0.1% Triton was applied for several washes. Samples were then incubated with the appropriate secondary antibodies conjugated to Alexa488, Alexa555 or Alexa647 fluorophores (1:500 to 1:1000 Molecular Probes) for 1h at RT. Cell nuclei were labeled using Hoechst 33342 (Life Technologies) to stain DNA. Immunolabeled samples were blinded upon mounting for subsequent imaging analysis.

### Immunohistochemistry

Immunohistochemistry was performed on postmortem motor cortex samples from three ALS patients carrying C9-HRE mutations and three non-neurological controls, obtained through the Northwestern University ALS Clinic, using previously described methods (Deng et al., 2011). Briefly, 6μm sections were cut from formalin-fixed and paraffin-embedded brain regions containing motor cortex. Sections were deparaffinized and rehydrated in serial solutions: 3 x 10 min in xylene, 3 x 5 min in 100% ethanol, 3 x 3 min in 95% ethanol, 1x 5 min in 75% ethanol, 1x 5 min in 50% ethanol, 1x 5 min in deionized water, and 1x 5 min in PBS. Antigen retrieval was performed using a decloaking chamber with Antigen Decloaker solution (Biocare Medical) at 125°C for 10 min. Sections were cooled to RT for 30 min and rinsed with deionized water. Samples were blocked with 1% BSA in PBS for 20 min at RT and subsequently incubated overnight at 4°C with primary antibodies: rRNA (mouse, 1:250, Novus Biologicals) and MAP2 (chicken, 1:500, Abcam). We next rinsed 3 x 5 min with PBS and incubated with appropriate secondary antibodies conjugated to Alexa488 or Alexa647 fluorophores (1:250, Molecular Probes) at RT for 45 min. Slides were rinsed 3 x 5 min and cell nuclei were labeled by DNA staining using Hoechst (Life Technologies) for 30 min at RT. To diminish autofluorescence, 0.3% sudan black in 70% ethanol was applied to each section for 45 sec and rinsed in deionized water for 5min. Slides were mounted in ProLong Diamond Antifade Mountant (Thermo Scientific) and labels were blinded for subsequent image analysis.

### *De novo* protein translation analysis

For single-cell protein translation analysis, we utilized a puromycin-based method termed SUnSET (Schmidt et al., 2009) that labels newly synthesized proteins. In short, cell cultures were pulsed for 5-10 min with puromycin (20 µM) at 37°C. Cells were then fixed and immunocytochemistry with anti-puromycin antibody was carried out as described above.

### Ribosomal RNA bait design

We design 20 nucleotide RNA baits based on the (GR)_50_-interacting 28S rRNA sequence identified by CLIP-Seq (Figure 3). We introduced 2’-O-methylations in the five nucleotides at the 5’ and 3’ ends to improve their stability (mGmGmUmCmUrCrCrArArGrGrUrGrArAmCmAmGmCmC). Using the same numbers of each of the 4 ribonucleotides we designed a scrambled sequence through the InvivoGen webtool https://www.invivogen.com/sirnawizard/scrambled.php (mGmCmAmCmUrGrArCrGrArUrGrCrGrCmUmAmGmCmA). RNA baits were delivered with Lipofectamine RNAiMAX transfection Reagent in Opti-MEM medium (Invitrogen) for 24 hours following manufacturer instructions. To evaluate cellular internalization of the RNA baits, we added a Cy3^TM^ fluorophore at the 3’ end of the ribonucleotide (mGmGmUmCmUrCrCrArArGrGrUrGrArAmCmAmGmCmC/3Cy3Sp/). All the RNA oligonucleotides utilized in this study were synthesized by Integrated DNA Technologies.

### Drosophila stocks

UAS-EGFP, UAS-FLAG-GR50-EGFP, and UAS-GR_36_ transgenic flies were generated as previously described for other *C9orf72* DPR fly models (Mizielinska et al., 2014; Wen et al., 2014). Fly stocks and crosses were maintained on standard fly medium in light/dark controlled incubators.

### *Drosophila* larval eclosion assay

UAS-EGFP and UAS-FLAG-GR50-EGFP lines were crossed with the motor neuron driver OK371-GAL4 on food mixed with and without 5 µM RNA Bait and incubated at 18 °C. 5 µM RNA Bait concentration was determined from (Zhang et al., 2015). The larvae were closely monitored from the 1st instar stage until they eclosed and become adults. The images of each developmental stage were taken using a Leica M205C dissection microscope equipped with a Leica DFC450 camera. Eclosion percentages were calculated as: (total number of eclosed adults) / (total number of pupal cases) x 100. Each condition was analyzed in triplicate using biological replicates.

### *Drosophila* eye degeneration assay

UAS-GR36 virgin females were crossed with GMR-GAL4 males on food mixed with and without 5 µM RNA Bait and incubated at 25 degrees. Adult eyes were imaged with a Leica M205C microscope 0-1 days post eclosion. Images of external eye phenotypes were then scored as previously described (Casci et al., 2019; Pandey et al., 2007). Briefly, eyes were examined for the presence or absence of the following features: supernumerary inter-ommatidial bristles, abnormal bristle orientation, ommatidial fusion, ommatidial pitting, disorganization of ommatidial array, and retinal collapse. If the following features were present, 1 point was given. Additional points were added corresponding to the total percentage of the effected eye area.

### *Drosophila* western blotting

Adult fly heads were collected from EGFP and FLAG-GR50-EGFP flies on RNA Bait food that eclosed and snap-frozen on dry ice. Heads were crushed on dry ice and incubated in RIPA buffer (150 mM NaCl, 1% NP40, 0.1% SDS, 1% sodium deoxycholate, 50 mM NaF, 2 mM EDTA, 2 mM DTT, 0.2 mM Na orthovanadate, 1 X Roche protease inhibitor #11836170001). Lysates were sonicated and centrifuged to remove debris. Supernatants were boiled in Laemmli buffer (Boston Bioproducts, #BP-111R) for 5 min and loaded onto 4%–12% Nupage Bis-Tris gels (Novex/Life Technologies). Proteins were transferred using the iBlot2 (Life Technologies, #13120134) onto nitrocellulose (iBlot 2 NC regular Stacks, Invitrogen, #IB23001). Western blots were blocked with milk solution (BLOT-QuickBlocker reagent, EMB Millipore, #WB57-175GM) and incubated with primary antibody overnight: chicken anti-GFP antibody, 1:3,000 (abcam) and mouse anti-tubulin, 1:10,000 (Sigma-Aldrich). Blots were washed and incubated in secondary antibody for one hour [anti-mouse IRDye 680D, 1:10,000 (LI-COR Biosciences); anti-Chicken IgY (H+L), 1:10,000 (ThermoFisher)] and imaged on a Licor imager (Odyssey CLx).

### *Drosophila* larval preparations and immunohistochemistry

Third instar larvae or adult *Drosophila* brains were dissected, fixed, and immunostained as previously described (Anderson et al., 2018). Briefly, animals were dissected in ice-cold phosphate buffered saline (PBS) (Lonza, #17–516 F), fixed in 4% formaldehyde, washed three times in PBS, incubated in 5% Triton X-100/PBS for 20 min, washed three times in 0.1% PBST (0.1 % Triton X-100/PBS), and incubated overnight with the primary antibody mouse anti-Lamin Dm0, 1:200 (Developmental Studies Hybridoma Bank). Larvae were washed three times in 0.1% PBST and incubated with secondary antibodies: anti-mouse Alexa Flour 647, 1:100 (Invitrogen, #28181. Stained larvae were mounted using DAPI Fluoroshield (Sigma-Aldrich, #F6182). Images were collected on a Nikon A1 eclipse T_i_ confocal microscope.

### Quantitative image acquisition and analysis

Images used for quantification were acquired at matched exposure times or laser settings and processed using identical settings. Quantifications were normalized within each respective experiment with n ≥ 3 independent experiments unless otherwise specified in figure legends. Image acquisition for HEK-293 experiments was performed on a Leica DMI4000B laser scanning confocal microscope (Leica, Buffalo Grove, IL) or with Leica DMi8 microscope (Leica, Buffalo Grove, IL) using a C10600-ORCA-R2 digital CCD camera (Hamamatsu Photonics, Japan), and processed with Fiji. For high-resolution images and 3D reconstructions Nikon W1 Dual CAM spinning Disk and Imaris Cell Imaging software were used.

Quantitative live cell imaging and analysis of GFP and GFP-(GR)x50 transduced iPSC-derived MNs was performed through an Incucyte System (Sartorious). Tracking of neuronal survival was manually performed with Fiji. Longitudinal tracking of iMNs was performed with a Molecular Devices ImageExpress once every 48 h. Neuronal survival was tracked using SVcell 3.0 (DRVision Technologies) or ImageJ. Neurons were scored as dead when their soma was no longer detectable by RFP fluorescence.

### Quantification and statistical analysis

All statistical analyses were done with Prism 7 software (GraphPad Software). Individual values were usually displayed by dots in the graphs, and represent all values measured in the study. The sample size (n) of each specific experiment are provided in the results section and the statistical test performed for each specific experiment is defined in the corresponding figure legend. For each statistical analysis, we first tested whether sample data fit into Gaussian distribution using the D’Agostino-Pearson omnibus normality test. To compare two experimental conditions, either a student’s t test (parametric) or a Mann-Whitney U test (non-parametric) was performed. To compare ≥ 3 experimental conditions, either a One-Way or Two-Way ANOVA followed by a Bonferroni post-hoc test (parametric) or a Kruskal-Wallis rank test followed by a two-stage linear step-up procedure of Benjamini, Krieger and Yekutieli (non-parametric) was performed. For comparisons of survival curves in Kaplan-Meier plots, we used Gehan-Breslow-Wilcoxon test.

## Notes

### Competing Interest Statement

The authors have declared no competing interest.

